# An intramolecular scrambling path controlled by a gatekeeper in Xkr8 phospholipid scramblase

**DOI:** 10.1101/2021.05.06.442885

**Authors:** Takaharu Sakuragi, Ryuta Kanai, Akihisa Tsutsumi, Hirotaka Narita, Eriko Onishi, Takuya Miyazaki, Takeshi Baba, Atsushi Nakagawa, Masahide Kikkawa, Chikashi Toyoshima, Shigekazu Nagata

## Abstract

Xkr8-Basigin is a phospholipid scramblase at plasma membranes that is activated by kinase or caspase. We investigated its structure at a resolution of 3.8Å. Its membrane-spanning region had a cuboid-like structure stabilized by salt bridges between hydrophilic residues in helices in the lipid layer. The molecule carried phosphatidylcholine in a cleft on the surface that may function as an entry site for phospholipids. Five charged residues placed from top to bottom inside the molecule were essential for providing a path for scrambling phospholipids. A tryptophan residue was present at the extracellular end of the pathway and its mutation made the Xkr8-Basigin complex constitutively active, indicating its function as a gatekeeper. The structure of Xkr8-Basigin provides novel insights into the molecular mechanisms underlying phospholipid scrambling.

## Introduction

The lipid bilayer in eukaryote plasma membranes comprises asymmetrically distributed phospholipids^1^. Phosphatidylserine (PtdSer) and phosphatidylethanolamine (PtdEtn) localize to the inner leaflet of the plasma membrane and phosphatidylcholine (PtdCho) to the outer leaflet. This asymmetrical distribution is disrupted in various biological processes, and PtdSer exposed to the cell surface activates enzymes or functions as a signal^2–4^. Activated platelets expose PtdSer as a scaffold for clotting enzymes, while PtdSer exposed on apoptotic cells functions as an “eat me” signal for macrophages.

The asymmetrical distribution of PtdSer and PtdEtn is maintained by ATP-dependent flippases (P4-ATPase), which translocate them from the outer to inner leaflets^5^. Flippases are inactivated in activated platelets or apoptotic cells^5,6^; however, this inactivation alone is insufficient for the quick exposure of PtdSer because it takes days for phospholipids carrying a hydrophilic head group to travel the hydrophobic lipid bilayer^7^. To expedite the exposure of PtdSer, cells carry scramblases that translocate phospholipids bidirectionally in an energy-independent manner^2,5^. We previously identified two membrane proteins (TMEM16F and Xkr8) as scramblases^8,9^ that are activated by distinct mechanisms. TMEM16F and several other TMEM16 family members, including its fugal homologues, function as Ca^2+^-dependent scramblases^8,10–14^. Xkr8 is a member of the XK family^9^ and forms a heterodimer with Basigin (BSG) or Neuroplastin (NPTN)^15^. It is activated by kinase or caspase to scramble phospholipids^9,16^. Xkr8 is responsible for exposing PtdSer in apoptotic cells, and its deficiency causes SLE-type autoimmune disease and male infertility^17,18^.

The tertiary structure showed that P4-ATPases and TMEM16F family members have hydrophilic clefts between peripheral transmembrane helices^12,19–22^, and supports an “out of the groove” model in which the phospholipid head group passes via the hydrophilic cleft while exposing its acyl chain to the lipid environment^23,24^. We herein report the structure of the human Xkr8-BSG heterodimer with a novel fold. It carries a hydrophobic cleft occupied by PtdCho on one side of the molecule and a hydrophilic path inside the molecule. A tryptophan residue at the extracellular end of the path appears to function as a gatekeeper, providing novel insights into the phospholipid scrambling mechanism.

## Results

### Purification of the hXkr8-hBSGΔ-Fab complex

The amino acid sequence of Xkr8 is well conserved in vertebrates (Extended Data Fig. 1). Xkr8 requires BSG or NPTN to localize to the plasma membrane^15^. We initially screened GFP-tagged Xkr8 orthologues for their expression and stability using fluorescence-detection size-exclusion chromatography, and found that human Xkr8 (hXkr8) and hBSG were relatively stable and abundantly expressed. BSG carries two immunoglobulin (Ig) domains, one of which is dispensable for its chaperone-like activity^15^. To enhance crystallization, we removed the first Ig and modified *N*-glycosylation sites (hBSGΔ) (Extended Data Fig. 2a). hBSGΔ supported the translocation of hXkr8 to the plasma membrane in *NPTN^-/-^BSG^-/-^*W3 (DKO)^15^ cells (Extended Data Fig. 2b), and transformed cells retained the ability to scramble phospholipids during apoptosis (Extended Data Fig. 2c).

hXkr8 was then fused to a histidine-tag and EGFP and co-expressed with hBSGΔ in Sf9 cells using a baculovirus system. The hXkr8-hBSGΔ complex purified from the lauryl-maltose neopentyl glycol (LMNG)-solubilized membrane fraction was nearly homogeneous (Extended Data Fig. 2d). Since the Fab of a monoclonal antibody often enhanced the crystallization of membrane proteins^25^, we prepared a Fab that preferentially recognized native over denatured hBSGΔ. X-ray crystallography of Fab and hXkr8-hBSGΔ-Fab revealed the structures of their extracellular regions at a high resolution of 2.51 Å (Extended Data Fig. 3 and Table S1). No structure of the transmembrane region may be probably because the transmembrane regions were lost during crystallization. To elucidate the structures of the transmembrane regions of hXkr8 and hBSG, free LMNG was removed by GraDeR^26^ (Extended Data Fig. 2e), and nearly homogeneous and monodisperse samples (Fig. 1a) were subjected to a cryo-EM single particle analysis.

**Fig. 1.**
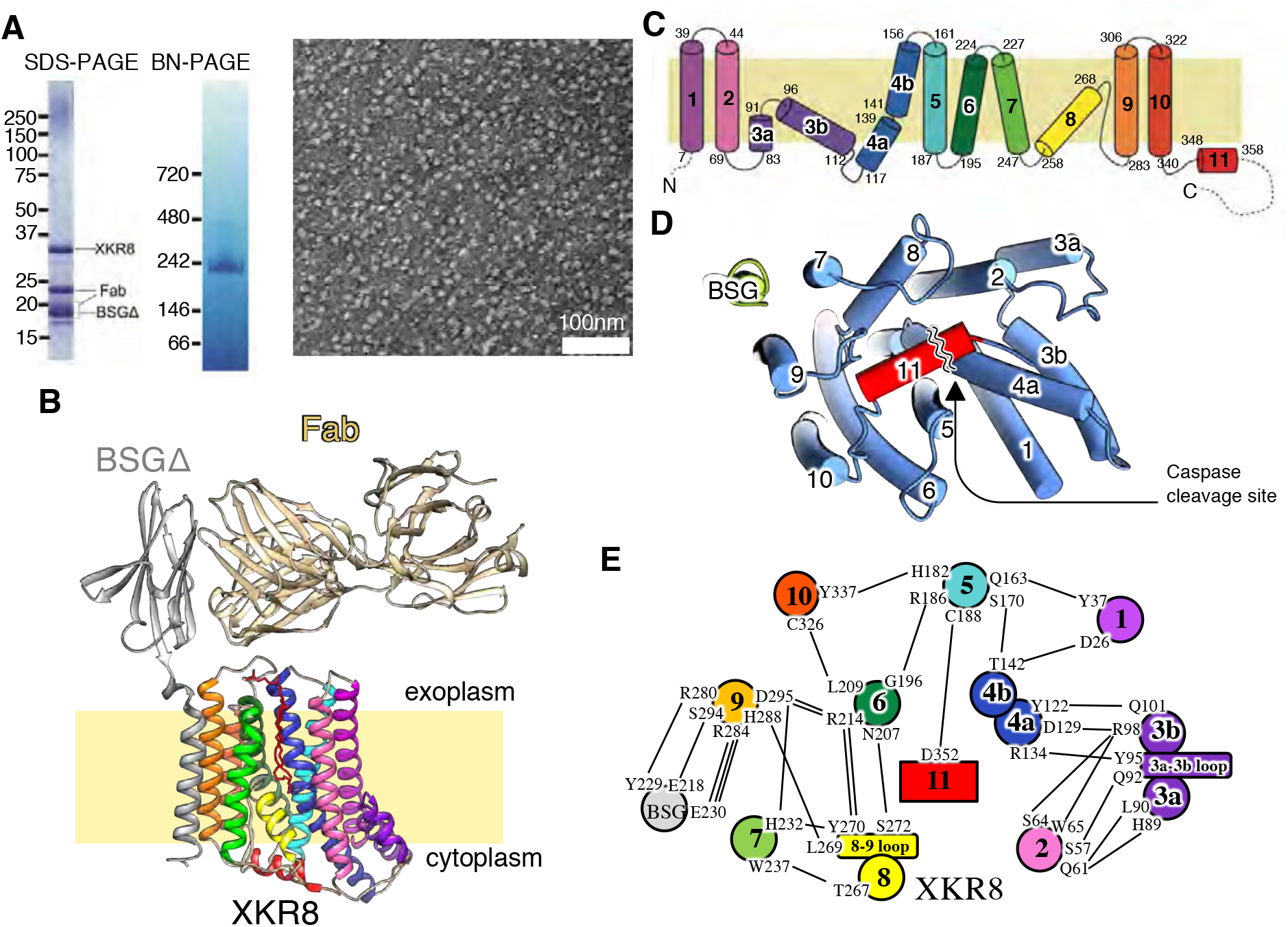
Structure of the hXkr8-hBSGΔ complex. **a,** Purification of the hXkr8-hBSGΔ-Fab complex. The purified hXkr8-hBSGΔ-Fab complex was analyzed by SDS-PAGE (3 μg), BN-PAGE (5 μg), or observed under an electron microscope (HITACHI H7650) after negative staining. **b**, The structure of the hXkr8-hBSGΔ-Fab complex. The location of the membrane is estimated from the position of tryptophan (Extended Data Fig. 6). **c**, α-Helices of hXkr8 are numbered and schematically shown. **d**, A view of the hXkr8-hBSGΔ complex from its cytoplasmic side. **e**, Amino acids connected via hydrogen bonds or a salt bridge in the hXkr8-hBSGΔ complex are shown.

### Structure of the hXkr8-hBSG complex

Despite the relatively small size of the hXkr8-hBSGΔ-Fab complex (110 kDa), electron micrographs showed well dispersed particles with sufficient contrast to reveal the structure of the hXkr8-hBSGΔ-Fab complex at a resolution of 3.8 Å (Extended Data Fig. 4 and Table S2). Based on the density map obtained by cryo-EM (Extended Data Fig. 5) and X-ray crystallography data, the structure of the hXkr8-hBSGΔ-Fab complex was elucidated (Fig. 1b). It consisted of hXkr8, hBSGΔ, and Fab at a 1:1:1 ratio, and PtdCho bound to the complex (Fig. 1b and Extended Data Fig. 5). The location of the membrane was tentatively assigned by referring to the position of tryptophan residues near the end of the helices^27,28^ (Extended Data Fig. 6). hXkr8 comprised 8 transmembrane helices (α1, α2, α4-7, α9, and α10), 2 helices traveling halfway to the membrane (α3 and α8), and one cytoplasmic helix (α11) (Fig. 1c). α11 contained a caspase 3-recognition sequence, interacted with α2, α4, α5, and α6 (Fig. 1d), and appeared to stabilize the disposition of these helices at the cytoplasmic side. The transmembrane region of hXkr8-hBSGΔhad a rectangular cuboid-like structure with a slight spread to the cytoplasmic face (Fig. 1b). Proteins with similar structures were not detected in a 3D homology search (PDBeFold).

### Interaction between hXkr8 and BSG

The structure of the hXkr8-hBSG complex as well as the analysis of hydrogen bonds and salt bridges indicated that α9 of hXkr8 was arranged near the transmembrane region of hBSGΔ (Fig. 1d and 1e). A207 and P211 of hBSGΔ were arranged toward T305 and T302 of hXkr8, respectively (Fig. 2a). To confirm their proximity, these residues were individually mutated to cysteine, tagged with GFP (for hXkr8) or HA (for BSG) (Fig. 2b), and expressed in HEK293T cells. Western blotting under non-reducing conditions showed a 100-kDa band with both anti-GFP and anti-HA when hXkr8-T305C was co-expressed with hBSG-A207C (Fig. 2c). When the membrane fraction was treated with the oxidant before SDS-PAGE, P211C of hBSGΔ and T302C of hXkr8 were also cross-linked (Fig. 2c), confirming the close proximity of these residues. At the cytoplasmic side of the hXkr8-hBSGΔ complex, E230 of hBSGΔ was close to Q247, R280, and R284 of hXkr8 (Fig. 2d). The E230A mutant of hBSG did not support the localization of hXkr8-GFP to the plasma membrane (Fig. 2e), while the R284E mutant of hXkr8 did not localize to the plasma membrane in PLB cells expressing intact hBSG (Fig. 2e), indicating that the interaction of E230 of BSG with R284 of hXkr8 was indispensable for the chaperone activity of BSG. An acidic residue (E218) in the middle of the transmembrane region of hBSG (Fig. 2f) was necessary for its chaperone activity for Xkr8^15^. The structure of the hXkr8-hBSGΔ complex indicated that E218 of hBSG interacted with S294 in α9 of hXkr8 (Fig. 2f). BSG, NPTN, and embigin (EMB) belong to the same family^15,29^. BSG and NPTN, but not EMB, have been shown to chaperone Xkr8 to the plasma membrane^15^. Using the *Coot* program^30^, residues in the transmembrane region of hBSG were replaced by the corresponding residues in hEMB (Fig. 2f). The replacement of G214 by valine resulted in a steric crash with L298 of hXkr8, and the replacement of I225 by threonine and Y229 by cysteine abolished their hydrophobic and polar interactions with I287 and R280, respectively. These results explain the inability of hEMB to chaperone Xkr8.

**Fig. 2.**
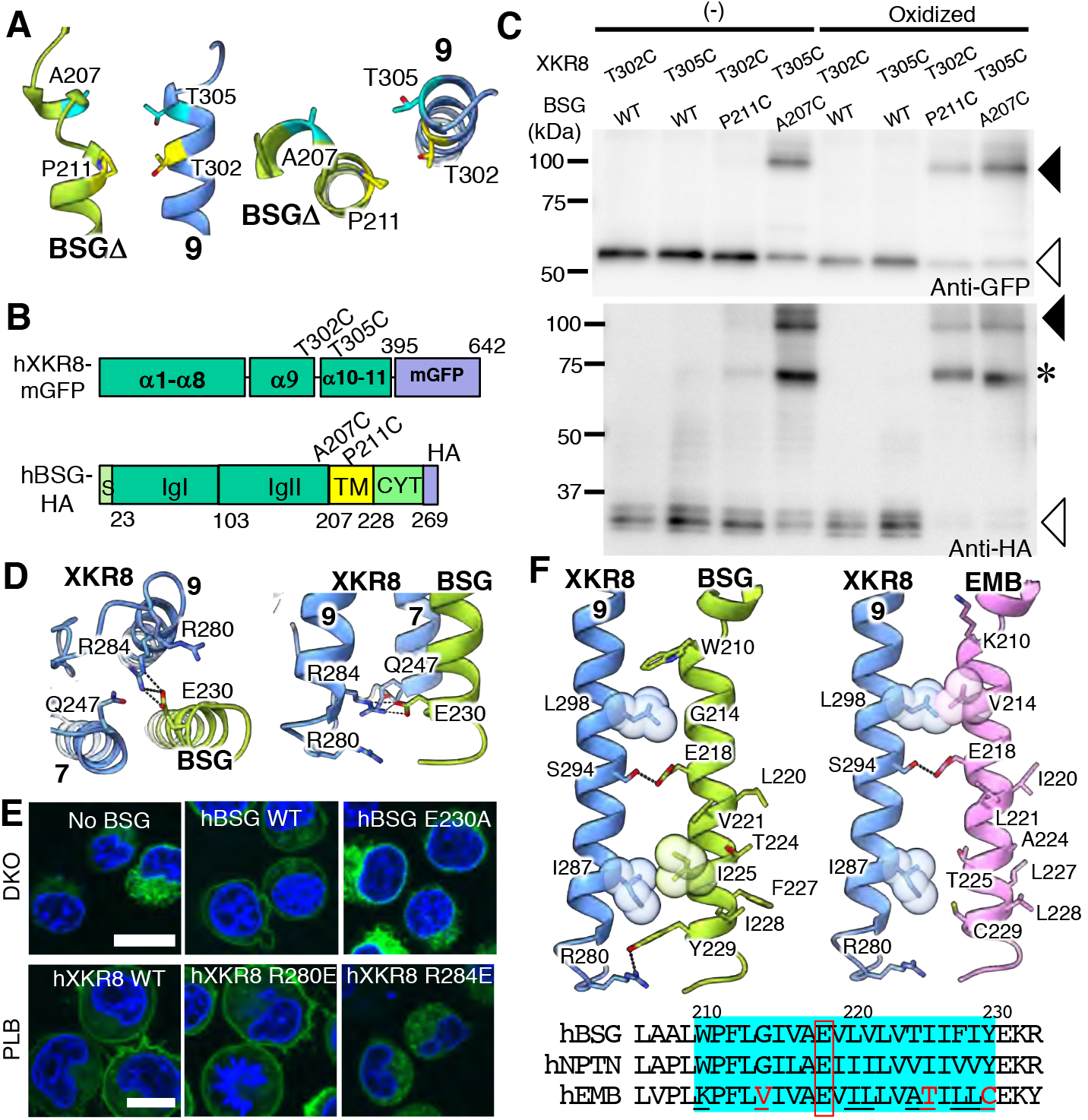
Interaction of hXkr8 with hBSGΔ. **a**, Side (left) and top (right) views of hXkr8-hBSGΔ near the extracellular surface. **b**, **c**, Cross-linking of cysteine-substituted hXkr8 and hBSG. The residues mutated to cysteine (T302 and T305 in hXkr8, and A207 and P211 in hBSG) are indicated in their schematic structure. In **c**, the indicated wild-type (WT) and mutant hXkr8-mGFP and hBSG-HA were expressed in HEK293T cells. Crude membrane fractions were treated or not with copper phenanthroline for oxidization, and subjected to Western blotting for GFP (upper panel) or HA (lower panel). Black arrowheads, cross-linked protein; white arrowheads, monomer. *, basigin dimer^41^. **d**, The interaction of hXkr8 with hBSGΔ near the cytoplasm is viewed from the cytoplasm or side. R284 of Xkr8 is connected with E230 of hBSG via hydrogen bonds. **e**, DKO expressing hXkr8-GFP with the wild-type or E230A-hBSG, or PLB expressing the EGFP-tagged wild-type, R280E-, or R284E-hXkr8 were observed for EGFP and Hoechst 33342. **f**, hXkr8-α9 was arranged with the transmembrane helix of hBSG or hEMB. Interacting residues are shown in the stick or sphere structure. A sequence alignment of the transmembrane region of hBSG, hNPTN, and hEMB is shown in the bottom.

### A hydrophilic pathway inside the molecule for scrambling phospholipids

The amino acid sequence of mouse (m)Xkr8 has 68.9% identity with hXkr8 (272/395 amino acids) (Fig. 3a), and homology modeling with the *Modeller* program indicated that the structure of mXkr8 was essentially identical to hXkr8 (Extended Data Fig. 7). An interesting feature of hXkr8 was the presence of 22 charged residues (Asp, Glu, Lys, and Arg) in the α-helices at the lipid layer (Fig. 3a). Sixteen of these residues were well conserved among vertebrates (Extended Data Fig. 1). To examine the role of these charged residues, we selected 11 residues (D12, D26, D30, R98, D129, E137, E141, D180, D295, R183, and R214) that were internally oriented (PDB). R214 and D295 were mutated to glycine or lysine, respectively, as found in XK in human patients with McLeod Syndrome^31,32^. Another 9 residues were mutated to Ala. The mutants were tagged by GFP at the C terminus and stably expressed in PLB985 or Ba/F3 cells. As shown in Fig. 3b, 4 mutants (R98A, D129A, R214G, and D295K) were not localized at the plasma membrane. A Western blot analysis showed that the expression of these mutant proteins was markedly weaker than that of wild-type Xkr8 (Fig. 3c). The structure of hXkr8 indicated two clusters of hydrophilic amino acids (one cluster with S64, R98, and D129, and another with R214, H232, Y270, and D295) in the transmembrane region of the complex (Fig. 3d). D129 in α4a formed salt bridges with R98 on α3b, while R214 on α6 and D295 on α9 formed a salt bridge. The weak expression of the mutant proteins (R98A, D129A, D295K, and R214G) indicated that the interaction between the hydrophilic residues in the transmembrane region plays an essential role in stabilizing the Xkr8-BSG complex.

**Fig. 3.**
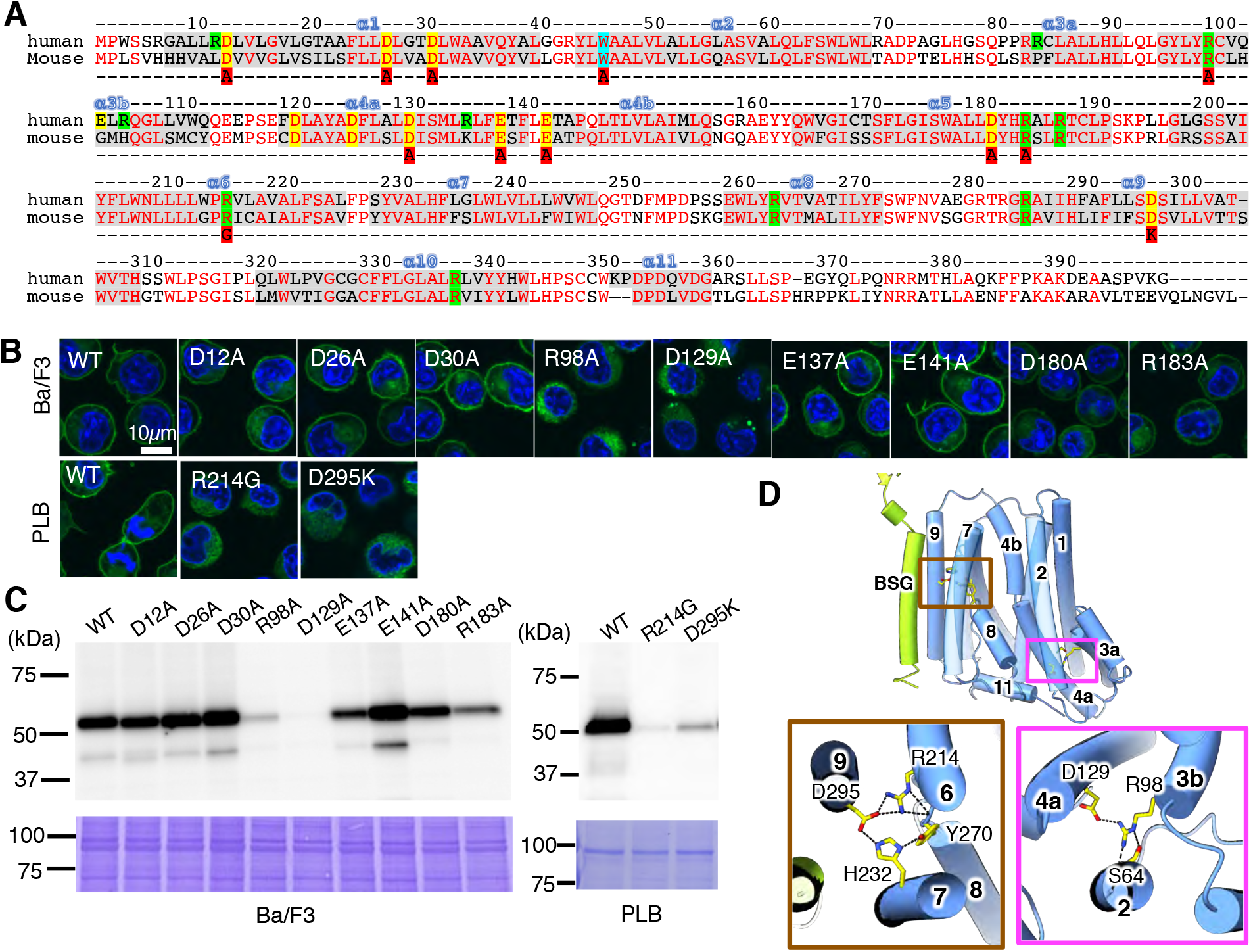
Charged residues in transmembrane helices of hXkr8. **a**, Sequences of hXkr8 (UniProt: Q9H6D3) and mXkr8 (UniProt: Q8C0T0) were analyzed by the MUSCLE Program (EMBL-EBI). Helices are shadowed and numbered. Conserved residues are in red. Negatively (Glu and Asp) and positively charged residues (Lys and Arg) in the lipid layer are highlighted in yellow and green, respectively, and Tryptophan-45 in light blue. The amino acids red-highlighted in the bottom line are mutated to Ala, Gly, or Lys. **b, c**, The GFP-tagged wild-type and indicated mutant mXkr8 (D12A, D26A, D30A, R98A, D129A, E137A, E141A, D180A, and R183A) or hXkr8 (R214G, and D295A) were stably expressed in mouse Ba/F3 or human PLB985, respectively, and observed under a fluorescent microscope (**b**). In **c**, whole cell lysates (Ba/F3) or the light membrane fraction (PLB) was analyzed by Western blotting with anti-GFP or stained by CBB. **d**, Stabilization of transmembrane α-helices by salt bridges. The regions at which helices α6-α9 and helices α4a and 3b interact with each other are expanded.

**Fig. 4.**
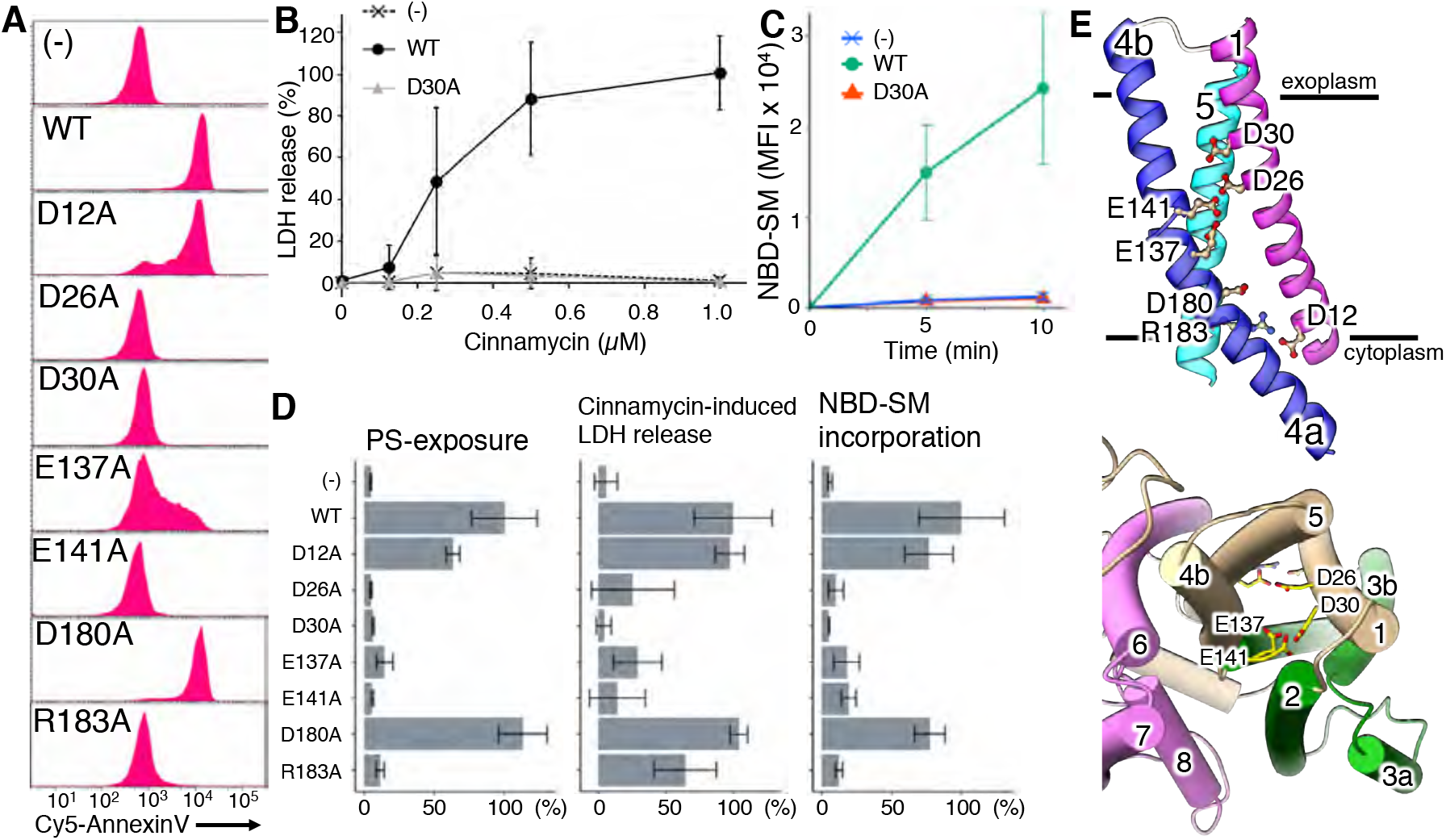
Requirement of charged residues in the membrane region of Xkr8 to scramble phospholipids. **a**, Ba/F3 transformants expressing the wild-type or indicated mutant mXkr8 were stained with Cy5-annexin V and analyzed by flow cytometry. **b**, Ba/F3 transformants expressing the wild-type or D30A mutant mXkr8 were incubated at 4°C for 15 min with the indicated concentration of cinnamycin and LDH released was expressed as a percentage of that released with 1% Triton-X100 (n=3) with S.D. **c**, Ba/F3 transformants expressing the wild-type or D30A mutant mXkr8 were incubated at 4°C with NBD-SM for the indicated periods. Incorporated NBD-SM was analyzed by flow cytometry, and its Mean Fluorescence Intensity (MFI) was plotted (n=3) with S.D. **d**, The scramblase activity of Ba/F3 transformants expressing the wild-type or indicated mutant mXkr8 was assayed for the exposure of PtdSer, sensitivity to cinnamycin-induced LDH release (0.5 μM for 15 min), and the incorporation of NBD-SM (5 min), as described above. Experiments were performed at least three times, and relative activities to that of the wild-type are shown with S.D. **e**, Close-up side and top views of charged amino acids (Glu, Asp, and Arg) in the α1, α4, and α5 helices. The side chains of charged amino acids are represented in a colored ball & stick model.

The 7 other mutants (D12A, D26A, D30A, E137A, E141A, D180A, and R183A) were expressed as efficiently as wild-type mXkr8 and localized at plasma membranes (Fig. 3b and c). The scrambling activity of mXkr8 was activated in mouse Ba/F3 cells by phosphorylation at the C-terminal tail region and was observed at 4°C, a temperature at which ATP-dependent flippase activity was reduced^16^. We used this system to examine the scrambling activities of mXkr8 mutants. As shown in Fig. 4a, Ba/F3 cell transformants expressing wild-type, D12, or D180 mutant mXkr8 constitutively exposed PtdSer. On the other hand, the exposure of PtdSer was markedly reduced in the transformants expressing D26A, D30A, E137A, E141A, and R183A mutants, indicating that these residues play an indispensable role in scrambling PtdSer from the inner to outer leaflets of plasma membranes. Xkr8 has been shown to non-specifically scramble phospholipids^9^. Accordingly, the treatment with cinnamycin (Ro 09-0918), which may kill cells exposing PtdEtn^33^, lysed BaF3 transformants expressing wild-type mXkr8, D12A, or D180A within 15 min (Fig. 4b and 4d). However, its killing activity was reduced in cells expressing D26A, D30A, E137A, E141A, or R183A mutants. Xkr8 scrambles phospholipids not only from the inner to outer leaflets, but also from the outer to inner leaflets^9^. Accordingly, wild-type mXkr8 as well as the D12A and D180A mutants of mXkr8 efficiently internalized NBD-SM (Fig. 4c and 4d). In contrast, this activity was markedly reduced by the mutation in D26, D30, E137, E141, and R183. The structure of hXkr8 indicated that these 5 amino acids were arranged from the top to bottom inside the molecule (Fig. 4e; PDB) and appeared to provide a path for scrambling phospholipids.

### A cleft for phospholipids on the surface and a gatekeeper for scrambling phospholipids

The hXkr8-hBSG complex contains a hydrophobic cleft facing the lipid bilayer in the upper middle part of the complex (Fig. 5a). The cleft was mainly generated by α2, α4, α6, and α7. Tilted α8 as well as the loop between α8 and α9 supported the cleft from below. An excess density that was consistent with PtdCho was detected in this cleft (Extended Data Fig. 5). Bound phospholipids were expected to originate from Sf9 cells because extra phospholipids were not added during protein preparation, indicating the strong affinity of hXkr8 for phospholipids. The lipid fit well in the cleft (Fig. 5a and 5b). The contact area between hXkr8 and PtdCho, calculated by Protein Interfaces, Surfaces and Assemblies (PISA)^34^, was approximately 800 Å^2^, which was similar to that observed between the PtdCho transfer protein (PCTP) and PtdCho (PDB ID: 1ln2)^35^. The two acyl chains of PtdCho in the narrow cleft were linked by more than 10 hydrophobic amino acid residues (Extended Data Fig. 8). The head group of PtdCho fit well into the space comprising R42, W45, Q155, and W309 (Fig. 5c). The arrangement of these amino acids with the lipid head group indicated that α2 and α4b were closely associated at their upper parts, and bound PtdCho appeared to contribute to stabilizing this association. This arrangement of amino acid residues also appeared to accept other phospholipids, such as PtdSer, and suggests that the cleft functions as the entry or exit point of the phospholipid scrambling path.

**Fig. 5.**
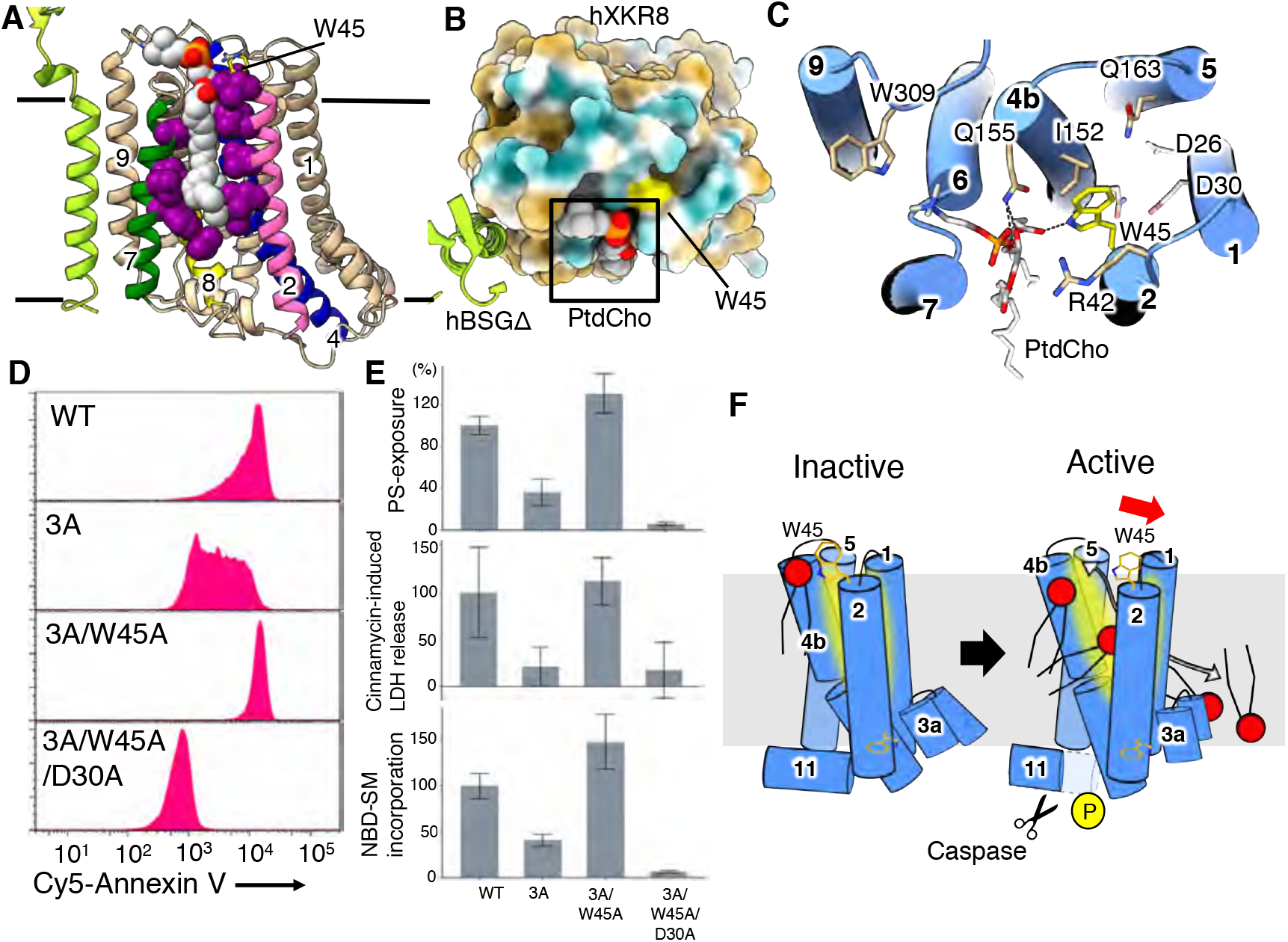
A lipid-binding pocket in hXkr8 and a gatekeeper for the phospholipid path. **a**, Front view of the hXkr8-hBSGΔcomplex. Hydrophobic residues surrounding PtdCho are in magenta spheres. PtdCho is in the silver sphere with the colored element. **b**, Top view of the complex. hXkr8 is in the colored surface view, brown, hydrophobic; blue, polar; white, neutral. PtdCho is boxed. **c**, A close-up top view of PtdCho with surrounding residues in hXkr8. The side chains of charged amino acids are represented in a stick model. The head group of PtdCho is coordinated with W309, Q155, W45, and R42. **d**, Ba/F3 cells were transformed with the wild-type, 3A (T356A/S361A/T375A), 3A/W45A, or 3A/W45A/D30A mutant of mXkr8-GFP, incubated with Cy5-Annexin V, and analyzed by flow cytometry. **e**, Scramblase activity in Ba/F3 transformants expressing the wild-type or indicated mutant mXkr8 was assessed for the exposure of PtdSer, sensitivity to cinnamycin-induced LDH release, and the incorporation of NBD-SM, as described in Fig. 4. Experiments were performed at least three times, and relative activities to that of the wild-type are shown with S.D. **f**, A potential activation mechanism of Xkr8. In the inactive state, W45 blocks phospholipids from entering the path comprised of charged residues (yellow-shaded). When Xkr8 is activated, W45 may tilt out to allow phospholipids to pass through the gate.

The cleft carrying PtdCho was detected at the front side of the scrambling path described above (Fig. 4e and 5c), and W45 localized at the top of the path appeared to prevent PtdCho from entering the path (Fig. 5c). To examine this possibility, W45 was mutated to alanine on the 3A mutant (T356A/S361A/T375A) of mXkr8, the scrambling activity of which was markedly reduced by the mutation of the three phosphorylation sites^16^ (Fig. 5d and 5e). On the other hand, the Ba/F3 transformants expressing the 3A-W45A mutant strongly exposed PtdSer, indicating that the W45A mutation provided mXkr8 constitutive-active or phosphorylation-independent scramblase. Similarly, sensitivity to the cytotoxicity of cinnamycin and the ability to incorporate NBD-SM were both reduced in the 3A mutant; however, this was rescued by the W45A mutation (Fig. 5e). The exposure of PtdSer, sensitivity to cinnamycin, and the incorporation of NBD-SM caused by the W45A mutation were completely blocked by mutating D30 to alanine (3A-D30A/W45A) (Fig. 5d and 5e). These results indicate that W45 serves as a gatekeeper for the path in mXkr8 to scramble phospholipids or serves as a key residue for transferring phospholipids in both the inward and outward directions at plasma membranes.

## Discussion

A critical step in the translocation of phospholipids between the outer and inner leaflets of lipid bilayers is the movement of their hydrophilic head group across the hydrophobic core, and scramblase may provide a hydrophilic path(s) to facilitate this process^36^. TMEM16F of Ca^2+^-dependent scramblases and its homologues exist as homodimers^8,10,12,13^, and an “out of the groove” model^23^ has been accepted for scrambling phospholipids; however, two different mechanisms have been proposed. One model suggests a Ca^2+^-induced conformational change to open the hydrophilic groove^19,37^, while another proposes lipid distortion and thinning to scramble phospholipids^38^.

The Xkr8-BSG complex has a cuboid-like structure with no similarities to the structures of TMEM16 family members. Several α-helices were connected by salt bridges between hydrophilic amino acids in the middle of transmembrane segments (BSG with α9, α6-9, and α4 with α3). A series of negatively or positively charged residues were placed on a “stairway” from the extracellular to intracellular border inside of the molecule. Mutational analyses indicated that at least 5 of them were indispensable for scrambling phospholipids, supporting their provision of a path for phospholipid scrambling. The mutation of 5 respective residues similarly affected the outward (PtdSer- and PtdEtn-exposure) and inward translocation (NBD-incorporation) of phospholipids, indicating that different phospholipids use the same path for inward and outward movements.

In the extracellular boundary of hXkr8 facing the lipid environment, we detected a hydrophobic cleft occupied by a single PtdCho molecule. The vestibule of this lipophilic cleft was near the extracellular end of the stairway, and a tryptophan (W45) localized between them (Fig. 5c). The replacement of W45 by alanine made Xkr8 scramblase constitutively active. In the resting state, tryptophan (W45) appears to function as a “gatekeeper” that prevents phospholipids from falling into the putative phospholipid path from the ectoplasmic side. Once phospholipids enter the path from the gate, they may automatically reach the inner leaflet. We previously postulated that the C-terminal tail region regulates or inhibits the scramblase activity of Xkr8^16^. In contrast, the constitutive exposure of PtdSer observed in the 3A-W45 mutant suggests that phospholipids freely enter the path from its cytoplasmic site in this mutant. Although we cannot rule out the possibility that the ability of the 3A mutant to restrict the entry of phospholipids from the cytoplasmic site is inhibited, the cleavage of the C-terminal region or its phosphorylation may cause a conformational change at the exoplasmic gate region.

Based on the present results, the following model was proposed the activation mechanism of Xkr8 to scramble phospholipids (Fig. 5f). Caspase 3 cleaves α11 or a putative kinase(s) phosphorylates the residues at the C-terminal tail region, which tilts α1-3 and α4a to open the W45 gate into the hydrophilic cluster. Phospholipids are recruited to the pathway by hydrophilic amino acids, slip down using them as “stepping stones”, and exit through crevasses between α1 and α5. Tryptophan (W176) and tyrosine (Y181) localized to the middle of α5 may destabilize the lipid bilayer and facilitate phospholipid scrambling^27^, as proposed for TMEM16F^38^. The XK family comprises 9 members in humans^39^, and its founding member (XK) is responsible for McLeod syndrome causing neuroacanthocytosis^40^. The two missense point mutations in XK identified in McLeod Syndrome corresponded to the pair of residues stabilizing the association of transmembrane helices in Xkr8, suggesting a similar structure for XK family members. The structure of the Xkr8-BSG complex elucidated herein will provide a template for understanding the molecular mechanisms underlying phospholipid scrambling and structure-function analyses of the XK family.

## Methods

### Cell lines, recombinant proteins, and materials

Spodoptera frugiperda (SF) 9 cells (ATCC CRL-1711) were cultured in Sf-900IIISFM (Gibco), human PLB985 (PLB) cells^1^ and *BSG^-/-^ NPTN^-/-^* W3 cells^2^ in RPMI containing 10% FCS and 50 μM β-mercaptoethanol, *TMEM16F^-/-^Xkr8^-/-^*Ba/F3^3^ in RPMI containing 10% FCS and 45 units/ml mouse IL-3, and human HEK293T cells in DMEM containing 10% FCS.

HRP-labeled rabbit anti-GFP Ab (anti-GFP pAb-HRP-DirecT) and HRP-labeled anti-HA Ab (Direct-Blot HRP anti-HA.11 Epitope Tag antibody) were obtained from MBL and BioLegend, respectively. Ro09-0198-biotin^4^ was a gift from Dr. Kohjiro Nagao (Graduate School of Engineering, Kyoto University). Cy5-labeled Annexin V was from BioVision. 1,2-dioleoyl-sn-glycero-3-phosphocholine (DOPC) was from Avanti Polar Lipids. (7S)-4-hydroxy-7-[(1R,2E)-1-hydroxy-2-hexadecen-1-yl]-N,N,N-trimethyl-14-[(7-nitro-2,1,3-benzoxadiazol-4-yl)amino]-9-oxo-3,5-dioxa-8-aza-4-phosphatetradecan-1-aminium, 4-oxide inner salt (NBD-SM) was from Cayman CHEMICAL. Lauryl maltose neopentyl glycol (LMNG), glycol-diosgenin (GDN), octaethyleneglycol mono-*n*-dodecylether (C_12_E_8_), and fluorinated fos-choline 8 (FC8) were from Anatrace. Cholesteryl hemisuccinate (CHS) and tobacco etch virus (TEV) protease were from Sigma-Aldrich.

### cDNAs and expression plasmids

DNA sequence coding for human Xkr8 (hXkr8) (UniProt: Q9H6D3) was custom-designed and synthesized by GeneArt (Thermo Fisher Scientific) to enhance mRNA stability and translational efficiency. hXkr8 cDNA was introduced into pBAC-3bV, a derivative of pFastBac1 (Thermo Fisher Scientific), designed to express the recombinant protein tagged C-terminally with a TEV cleavage sequence, 8 histidine residues, a c-Myc, 8 histidine residues, and monomeric (m)EGFP (A206K) in this order^5^. This construct (pBAC-3bv carrying hXkr8) was hereafter referred to as hXkr8-TH-mGFP. cDNA for human basigin (hBSG) (UniProt: P35613-2) was described previously^2^. hBSG cDNA lacking the Ig-like domain (Ig) 1 (aa 23-102) was produced by PCR, fused to a FLAG epitope at the C terminus, and introduced into pBAC-3bV vector using In-Fusion Cloning Kits (Takara Bio). Two N-glycosylation sites (NGS and NGT) in the Ig2 domain at positions 152 and 186 were mutated to QGS and QGT by PCR, and designated as hBSGΔ. Primers used for PCR to produce non-glycosylated mutant hBSG cDNA were as follows. N152Q, 5’-CTCATGCAAGGCTCCGAGAGCAGGTTCTTC-3’ and 5’-GGAGCCTTGCATGAGGGCCTTGTCCTCAGAG-3’; N186Q, 5’-CGGTGCCAAGGCACCAGCTCCAAGGGC-3’ and 5’-GGTGCCTTGGCACCGGTACTGGCCGGG-3’.

cDNAs for mouse BSG (mBSG), mouse Xkr8 (mXkr8), and mXkr8 S/T-3A (T356A/S361A/T375A) mutant were described previously^2,3^. Primers for mXkr8 mutants (D12A, D26A, D30A, R98A, D129A, E137A, E141A, D180A, R183A, W45A), hXkr8 mutants (R134A, R214G, D295K, R280E, R284E, T302C, and T305C) and hBSG mutants (A207C, P211C, and E230A) were as follows. mXkr8 D12A; 5’-GCCTTAGCCGTGGTCGTAGGCCTGGTG-3’ and 5’-GACCACGGCTAAGGCCACATGGTGGTGC-3’, mXkr8 D26A; 5’-CTGCTGGCTCTGGTCGCTGACCTGTGG-3’ and 5’-GACCAGAGCCAGCAGGAAAGACAAGATACTCAC-3’, mXkr8 D30A; 5’-GTCGCTGCCCTGTGGGCCGTTGTCCAG-3’ and 5’-CCACAGGGCAGCGACCAGATCCAGCAGG-3’, mXkr8 R98A; 5’-CTGTATGCGTGTTTGCACGGAATGCATCAAGG-3’ and 5’-CAAACACGCATACAGGTAGCCGAGCTGCAG-3’, mXkr8 D129A; 5’-TCCCTGGCCATCAGCATGCTGAAGCTTTTCGAG-3’ and 5’-GCTGATGGCCAGGGAGAGAAAGTCTGCGTAG-3’, mXkr8 E137A; 5’-CTTTTCGCGAGCTTCCTGGAGGCGACG-3’ and 5’-GAAGCTCGCGAAAAGCTTCAGCATGCTGATG-3’, mXkr8 E141A; 5’-TTCCTGGCGGCGACGCCACAGCTC-3’ and 5’-CGTCGCCGCCAGGAAGCTCTCGAAAAGCTTC-3’, mXkr8 D180A; 5’-CTGCTGGCTTACCATCGGTCTCTGCGTACC-3’ and 5’-ATGGTAAGCCAGCAGTGCCCACGAGATG-3’, mXkr8 R183A; 5’-TACCATGCGTCTCTGCGTACCTGTCTTCCC-3’ and 5’-CAGAGACGCATGGTAATCCAGCAGTGCCCAC-3’, mXkr8 W45A; 5’-TTATCTGGCGGCCGCGCTGGTACTGG-3’ and 5’-GCGGCCGCCAGATAACGGCCAAGGAGCACG-3’, hXkr8 R134A; 5’-ATGCTGGCCCTGTTCGAGACATTCCTGGAAACCG-3’ and 5’-GAACAGGGCCAGCATGGAGATGTCCAGGGC-3’, hXkr8 R214G; 5’-GTGGCCCGGAGTGCTGGCCGTGGCC-3’ and 5’-AGCACTCCGGGCCACAGCAGCAGC-3’, hXkr8 D295K; 5’-CTGAGCAAGAGCATCCTGCTGGTGGCTAC-3’ and 5’-GATGCTCTTGCTCAGCAGAAAAGCGAAGTGG-3’, hXkr8 R280E; 5’-AGGGCGAGACAAGAGGCAGAGCCATCATC-3’ and 5’-CTCTTGTCTCGCCCTCGGCCACGTTG-3’, hXkr8 R284E; 5’-AGAGGCGAAGCCATCATCCACTTCGCTTTTC-3’ and 5’-GATGGCTTCGCCTCTTGTCCGGCCCTC-3’, hXkr8 T302C; 5’-GTGGCTTGCTGGGTCACCCACTCTAGCTGG-3’ and 5’-GACCCAGCAAGCCACCAGCAGGATGCTGTC-3’, hXkr8 T305C; 5’-TGGGTCTGCCACTCTAGCTGGCTGCCTAGC-3’ and 5’-AGAGTGGCAGACCCAGGTAGCCACCAGCAG-3’, hBSG A207C; 5’-CACCTGTGCGCCCTCTGGCCCTTCCTGG-3’ and 5’-GAGGGCGCACAGGTGGCTGCGCACGC-3’, hBSG P211C; 5’-CTCTGGTGCTTCCTGGGCATCGTGGCTGAGG-3’ and 5’-CAGGAAGCACCAGAGGGCGGCCAGGTG-3’, hBSG E230A; 5’-ATCTACGCGAAGCGCCGGAAGCCC-3’ and 5’-GCGCTTCGCGTAGATGAAGATGATGGTGACCAGC-3’. The resultant cDNAs were introduced using In Fusion HD Cloning Kits into pMXs puro c-GFP^1^, pCX4-bsr c-HA^2^, pPEF-mEGFP-FLAG^3^, or pNEF-HA, a derivative of pEF-BOS-EX^6^ carrying a neomycin resistance gene and DNA fragment for a HA tag. The authenticity of cDNAs was confirmed by DNA sequencing.

### Purification of the hXkr8-hBSGΔ complex

Recombinant baculovirus carrying hXkr8-TH-mGFP or hBSGΔ cDNA was obtained with the Bac-to-Bac Baculovirus Expression System (Thermo Fisher Scientific) according to the manufacturer’s instructions. Sf9 cells (4 × 10^6^ cells/ml) were co-infected with baculoviruses carrying hXkr8-TH-mGFP or hBSGΔ for large-scale expression. At 60-h post-infection, cells were harvested by centrifugation (1,000 × *g* for 3 min).

In cryo-EM grid preparation, the hXkr8-hBSGΔ complex was purified as follows. Infected Sf9 cells (36 g) were suspended in 150 ml of 20 mM HEPES-NaOH buffer (pH 7.0) containing 150 mM NaCl, 1 mM EDTA, 1 mM EGTA, 0.4 mM Pefabloc SC, 1 mM pAPMSF, 1 mM Tris(2-carboxyethyl)phosphine (TCEP), and a protease inhibitor mixture (cOmplete, EDTA-free, Roche Diagnostics). After adding benzonase (Merck) to a final concentration of 1.3 units/ml, cells were disrupted on ice by sonication using an ultrasonic disrupter (Sonifier 450 Advanced; Branson) (0.5-sec sonication at an output level of 4 and 0.5-sec cooling intervals for a total sonication period of 2.5 min). Unbroken cells and nuclei were removed by centrifugation at 6,000 × *g* for 25 min, and crude membrane fractions were collected by centrifugation at 99,650 × *g* for 1 h using a Beckman TYPE 70 Ti rotor.

Membranes were homogenized using a Dounce homogenizer with a tightly fitting pestle in 80 ml of 20 mM HEPES-NaOH buffer (pH 7.0) containing 150 mM NaCl, 25% glycerol, 50 mM imidazole, 0.4 mM Pefabloc SC, 1 mM pAPMSF, 1 mM TCEP, and cOmplete EDTA-free. Homogenates were subjected to sonication (0.5-sec flash with 0.5-sec cooling; total sonication time of 30 sec) at an output level of 3 using Sonifier 450 Advanced. After the addition of LMNG and CHS to final concentrations of 1.0 and 0.1%, respectively, the sample was incubated at 4°C for 2 h and centrifuged at 99,650 × *g* for 1 h to remove insoluble materials. The lysates (570 mg protein) were loaded onto a 5.0-ml HisTrap High Performance column (Cytiva) attached to high-performance liquid chromatography system (HPLC, Prominence; Shimadzu), and washed with 25 ml of increasing concentrations (60-80 mM) of imidazole in Buffer A [20 mM HEPES-NaOH buffer (pH 7.0), 0.01% LMNG, 0.001% CHS, 150 mM NaCl, and 1mM TCEP] containing 25% glycerol. The hXkr8-TH-mGFP/hBSGΔcomplex was then eluted with 45 ml of a linear gradient of the imidazole concentration from 80 to 400 mM in Buffer A containing 25% glycerol. Fractions carrying the hXkr8-TH-mGFP/hBSGΔ complex were pooled and treated at 4°C with 1,600 units of TEV protease for 12-18 h while dialyzing against Buffer A containing 10% glycerol and 1 mM EDTA. After confirming the cleavage by SDS-PAGE, the sample was re-loaded over the HisTrap column equilibrated with 20 mM HEPES-NaOH buffer (pH 7.0), 0.005% LMNG, 0.0005% CHS, 150 mM NaCl, 10% glycerol, and 1 mM TCEP to trap His-tagged GFP and TEV protease. Fractions flowed through the column were collected and concentrated approximately 100-fold by ultrafiltration at 5,000 × g using an Amicon Ultra-15 50K filter.

The hXkr8-hBSGΔ complex for crystallization was prepared using a similar procedure. In brief, baculovirus-infected Sf9 cells (27 g) were suspended in 100 ml of TBS [20 mM Tris-HCl (pH 7.5) and 150 mM NaCl] containing 1 mM EDTA, 1 mM EGTA, 0.4 mM Pefabloc SC, 1 mM DTT, 1.3 units/ml benzonase, and cOmplete EDTA-free. Cells were disrupted by sonication, and membrane fractions were collected by centrifugation as described above. Membranes were then homogenized using a Dounce homogenizer in 60 ml of TBS containing 25% glycerol, 50 mM imidazole, 0.4 mM Pefabloc SC, and cOmplete EDTA-free, which was followed by sonication and solubilization with 1% LMNG and 0.1% CHS as described above. The hXkr8-TH-mGFP/hBSGΔ complex was purified by the HisTrap column as described above, except that TBS-based buffer (pH 7.5) was used instead of HEPES-based buffer (pH 7.0). After eluting the hXkr8-TH-mGFP/hBSGΔ complex from the column with a linear gradient of imidazole concentrations (80 to 400 mM), the sample (2 mg hXkr8-TH-mGFP /hBSGΔ protein) was treated at 4°C with 1,200 units of TEV protease, while sequentially dialyzing against TBS containing 0.01% LMNG, 0.001% CHS, 10% glycerol, 1 mM DTT, 1 mM EDTA, and 1 mM EGTA for 12-14 h, and against TBS containing 0.01% LMNG, 0.001% CHS, and 10% glycerol for 3-4 h. GFP and TEV protease were removed by the His-Trap column in TBS containing 0.005% LMNG and 10% glycerol, and proteins that were not attached to the column were concentrated as described above.

### Monoclonal antibodies against the hXkr8-hBSG complex

Rabbit monoclonal antibodies against the native hXkr8-hBSGΔ complex were established as previously described^7^. In brief, 4 New Zealand White (NZW) rabbits from KITAYAMA LABES Co., LTD. were immunized with the purified hXkr8-hBSGΔ complex using TiterMax^TM^ or aluminum salts as adjuvants. IgG-positive B cells were sorted from peripheral mononuclear cells and splenocytes, and cultured at 37°C for 11 days at 2 cells/well in 96-well round-bottomed plates together with 2.5 × 10^4^ Mitomycin C-treated murine EL-4 feeder cells. A total of 14,080 clones were screened for the ability to recognize the hXkr8-hBSGΔ complex using the Octet Red96 System (ForteBio), and two clones (XBA14 and XBA18) were positive. The amino acid sequences of the variable regions of the XBA14 and XBA18 antibodies differed at 4 positions (2 in heavy chains and 2 in light chains). However, the CDR3 domain of their heavy chains had the same sequences, suggesting that these antibodies recognize the same epitope. The DNA segments coding for the variable region of their heavy and light chains were inserted into the pEF-BOS-based expression vector carrying the constant region of human IgG heavy or light chain. The recombinant IgG (XBA14 and XBA18) was produced in FreeStyle 293 cells (Thermo Fisher Scientific), and purified by Protein A-Sepharose (Cytiva) or MabSelect SuRe (Cytiva).

All animal care and experimental protocols were performed in accordance with the guidelines for the care and use of laboratory animals at Chugai Pharmaceutical Co., Ltd. The protocol was approved by the Institutional Animal Care and Use Committee at Chugai Pharmaceutical Co., Ltd.

### Preparation of the hXkr8-hBSGΔ-Fab complex

To prepare the Fab fragment of XBA mAb, 15 mg IgG was digested at 35°C for 2 h with 2 μg Lys-C in Tris-HCl buffer (pH 8.0), and Fc fragments were removed using HiTrap SP HP (Cytiva) and MabSelect SuRe columns. Fab fragments (Fab14 and Fab18) were purified by gel filtration (Superdex 200 pg 16/600, Cytiva) in 20 mM HEPES-NaOH (pH 7.1) containing 100 mM NaCl.

To prepare the hXkr8-hBSGΔ-Fab complex, 1.1 mg of hXkr8-hBSGΔ was incubated with 3.6 mg of Fab14 at 4°C for 20-30 h in 0.48 ml of 0.36-0.004% LMNG, 16 mM Tris-HCl (pH 7.5), 4 mM HEPES-NaOH (pH 7.1), 140 mM NaCl, and 8% glycerol. The sample was then subjected to gel filtration using the Superose 6 Increase 10/300 GL column (Cytiva) equilibrated with TBS containing 10% glycerol. Fractions containing hXkr8-hBSGΔ-Fab were collected and concentrated by ultrafiltration using Vivaspin 2-100K (Cytiva).

In the cryo-EM analysis, detergents in the sample were removed by GraDeR as described^8^. In brief, 0.9 mg of hXkr8-hBSGΔ was incubated with 1.8 mg of Fab18 at 4°C for 20-30 h in 0.36 ml of 20 mM HEPES-NaOH buffer (pH 7.0), 0.004-0.36% LMNG, 0.0004-0.036% CHS, 136 mM NaCl, 7% glycerol, and 0.7 mM TCEP. Using a Gradient Master (BIO COMP), a linear gradient of glycerol (10 to 30%) with a reverse gradient of detergents (0.00225 to 0% for LNMG and 0.00075 to 0% for GDN) was established in 4 ml of 20 mM HEPES-NaOH buffer (pH 7.0) containing 100 mM NaCl, 1 mM EGTA, 1 mM TCEP, 10 µM *p*APMSF, and 100 µM Pefabloc SC. Aliquots (0.1-0.2 ml) of protein samples were loaded on the top of the gradient, and centrifuged at 200,614 × *g* for 15 h using a Beckman SW55Ti rotor. Gradients were fractionated using the Piston Gradient Fractionator (BIO COMP), and fractions carrying hXkr8-hBSGΔ-Fab were pooled and concentrated by ultrafiltration using Vivaspin 2-100K. Glycerol was removed by repeating concentration and 30-40-fold dilution with the grid buffer [20 mM HEPES-NaOH (pH 7.0), 100 mM NaCl, 1 mM EGTA, 1 mM TCEP, 10 µM *p*APMSF, and 100 µM Pefabloc SC] three times.

### Crystallization and structural elucidation of the hXkr8-hBSGΔ-Fab14 complex

Fab14 was crystallized using the sitting drop vapor diffusion technique in the initial screening with PEGRx 2 primary screen variables (Hampton), and its structure was elucidated by an X-ray diffraction analysis. In brief, 100 nl of 7.1 mg/ml Fab14 was mixed with an equal volume of reservoir solution (0.1% n-Octyl-β-D-glucoside, 0.1 M sodium citrate tribasic dihydrate, and 22% PEG3350) and kept at 20°C. Crystals were harvested after 10 days, cryoprotected by soaking in 20 mM HEPES-NaOH buffer (pH 7.1) containing 0.1% n-Octyl-β-D-glucoside, 0.1 M sodium citrate tribasic dihydrate, 22% PEG3350, 100 mM NaCl, and 15% ethylene glycol, and flash-frozen in liquid nitrogen. Diffraction data were collected by helical scanning at BL32XU (SPring-8) on an EIGER X 9M detector (Dectris) and processed with autoPROC^9^, STRANISO^10^, and the CCP4 program^11^. The structure was elucidated by molecular replacement with Molrep^12^ using the coordinate of the anti-hinge rabbit antibody (PDB 4ma3) as a search model. The model was refined by Refmac5^13^, phenix.refine^14^ and *Coot*^15^. The final model for Fab14 included all amino acid residues, except for residues from positions 127 to 132 of the heavy chain because they were not visible in the electron density map.

The crystallization of hXkr8-hBSGΔ-Fab14 was also performed by the sitting drop vapor diffusion method with the MemGold^TM^ screening system (Molecular Dimensions). In brief, 30 µl of 8.1 mg/ml of hXkr8-hBSGΔ-Fab14 in a glass tube coated with a film of 90 µg DOPC was supplemented with 225 µg of C_12_E_8_ (lipid detergent ratio of 1:2.5) and incubated overnight while mixing. One hundred-nanoliter aliquots of the lipidated protein were mixed with an equal volume of 0.1 M Tris-HCl (pH 8.0) buffer containing 0.1 M sodium chloride, 0.1 M cadmium chloride hemi(pentahydrate), and 33% PEG400, and kept at 20°C. Crystals were harvested after 2 months and flash-frozen in liquid nitrogen. X-ray diffraction data were collected by helical scanning at the BL32XU beamline at SPring-8 and processed with autoPROC^9^, STARANISO^10^, and CCP4^11^ program. The phase was identified by molecular replacement by Molrep^12^ using a coordinate of Fab14 as a search model, and domain 2 of Basigin (PDB:3B5H) was manually placed by the *Coot* program^15^. The model was refined by the *Coot*^15^ and phenix.refine^14^ programs, and the final model revealed the structure of Fab14 and extracellular region of hBSGΔ. Other parts including hXkr8 and the transmembrane region of hBSG were not modeled because they were not visualized in the electron density map.

### Electron microscopy

In negative-staining electron microscopy, a 3-μl aliquot of the hXkr8-hBSGΔ-Fab complex (0.01 mg/ml) purified through GraDeR was applied to freshly glow-discharged, carbon-coated 200 mesh copper grids (Nisshin EM). After briefly blotting with filter paper (Whatman #1), samples were stained with 2% uranyl acetate, blotted again, and subjected to transmission electron microscopy (H7650, HITACHI) at the Center for Medical Research and Education, Osaka University.

In cryo-EM, the hXkr8-hBSGΔ-Fab complex purified through the GraDeR procedure was adjusted to 8 mg/ml with the grid buffer, and fluorinated FC8 was added at a final concentration of 0.075%. An aliquot (3 µl) of the purified protein was applied to the glow-discharged Quantifoil holey carbon grid (Cu/Rh, R1.2/1.3, 200 mesh), blotted by Vitrobot (Thermo Fisher Scientific) for 4 s under 100% humidity at 6°C with a blot force of +10, and plunge-frozen in liquid ethane. Movies were acquired by Titan Krios (Thermo Fisher Scientific) equipped with the K3 summit direct electron detector (Gatan) operated at 300 kV working under low-dose conditions (48 frames at 48 *e^-^* Å^-2^).

Movies were subjected to beam-induced motion correction using RELION3.1, and contrast transfer function (CTF) parameters were estimated by CTFFIND4^16^. All of the following processes were performed using RELION 3.1. To generate 2D templates for automatic particle picking, particles were picked from 200 randomly selected micrographs using template-free Laplacian-of-Gaussian picking, then subjected to multiple rounds of reference-free 2D classification. Good 2D classes were selected as templates and 2D template-based particle picking was performed. In total, 864,702 particles from 1,977 micrographs were auto-picked and extracted with down-sampling to a pixel size of 3.32 Å/pix. Particles were subjected to 2D classification. The 655,655 particles selected were subjected to 3D classification and 192,162 particles were selected, re-extracted with a pixel size of 1.245 Å/pix, and subjected to 3D refinement. The 3D map and particle set yielded were subjected to per-particle defocus refinement, beam-tilt refinement, Bayesian polishing, and 3D refinement. No-align focused 3D classification with a mask covering hXkr8-hBSGΔ and the Fv domain of Fab resulted in one class of map with 62,124 particles. Final 3D refinement and postprocessing yielded a map with an overall resolution of 3.8 Å, estimated by the gold-standard FSC = 0.143 criteria. The processing strategy is described in fig. S4.

### Model building and refinement

At a resolution of 3.8 Å, the cryo-EM density map had sufficient quality for the *de novo* model building of hXkr8 and transmembrane helix of hBSG. A polyalanine helix model was initially built by *Coot*^15^, and amino acids were subsequently assigned based on the density of bulky side chains. The structure of the extracellular domain of hBSG and Fab, elucidated by the X-ray diffraction analysis, was fit to the map by rigid body refinement (phenix.real_space_refine)^17^. The model was then manually fit to the map using *Coot*^15^, and subjected to simulated annealing at 500K using the program of phenix.real_space_refine. The first 6 amino acids at the N terminus and last 35 amino acids at the C terminus of hXkr8 as well as the last 35 amino acids in the C terminus of hBSG were not modeled. Molecular graphics were illustrated using UCSF Chimera^18^ and UCSF ChimeraX^19^.

### Expression level of Xkr8

Cell lysates were prepared by incubating cells on ice for 10 min in radioimmunoprecipitation assay (RIPA) buffer [50 mM Hepes-NaOH buffer (pH 8.0), 1% Nonidet P-40, 0.1% SDS, 0.5% sodium deoxycholate, and 150 mM NaCl] containing c0mplete EDTA-free. After removing insoluble materials by centrifugation at 20,000 × *g* at 4°C for 10 min, the lysates were mixed with a 0.25 volume of 5 × SDS sample buffer [200 mM Tris-HCl buffer (pH 6.8), 10% SDS, 25% glycerol, 5% 2-mercaptoethanol, and 0.05% bromophenol blue]. Samples were incubated at room temperature for 30 min, separated on 10-20% SDS-PAGE (Nacalai Tesque), transferred to a PVDF membrane (Millipore), blotted with HRP-labeled rabbit anti-GFP Ab (anti-GFP pAb-HRP-DirecT), and visualized using Immobilon Western Chemiluminescent HRP substrate (Merck Millipore).

In some cases, the expression of Xkr8 was examined with the light membrane fractions as previously described^2^. Briefly, cells from the 200-ml culture were homogenized with a Dounce homogenizer in 2.5 ml of 10 mM Tris-HCl buffer (pH 7.5) containing 1 mM *p*APMSF and mixed with 2.5 ml of 10 mM Tris-HCl buffer (pH 7.5), 0.5 M sucrose, 0.1 M KCl, 10 mM MgCl_2_, 1 mM EGTA, and 1 mM *p*APMSF. After successive centrifugation at 800 × *g* for 10 min and at 8,000 × *g* for 10 min, membrane fractions were collected by centrifugation at 100,141 × *g* for 1 h using the Beckman SW55Ti rotor. Crude membranes were suspended in centrifugation buffer [10 mM Tris-HCl buffer (pH 7.5), 50 mM KCl, 5 mM MgCl_2_, 0.5 mM EGTA, and 1 mM *p*APMSF] containing 40% (w/v) sucrose and loaded between 17% (w/v) and 50% (w/v) sucrose layers in centrifugation buffer. After centrifugation at 100,141 × *g* at 4°C for 150 min, the light membrane fraction at the 17% (w/v) /40% (w/v) sucrose interface was collected with a 21G needle. The sample was diluted with centrifugation buffer to reduce the sucrose concentration to 3-4%, and subjected to centrifugation at 100,141 × *g* for 1 h. After homogenizing the precipitates in 100 µl of TBS containing 10% glycerol and c0mplete EDTA-free protease inhibitors by passing through a non-dead-space 29G needle (Myjector; TERUMO), a 0.1 volume of 10% LMNG −1% CHS was added to the sample and rotated at 4°C for 2 h for solubilization. Insoluble materials were then removed by spinning at 104,300 × *g* at 4°C for 10 min using the Beckman TLA100.4 rotor. The supernatants were recovered as light membrane lysates, and subjected to Western blotting as described above.

### Cross-linking experiment

The cross-linking of Cys-substituted hXkr8 and hBSG was performed as described^20^. Briefly, HEK293T cells were co-transfected with pEF-BOS-based expression vectors for hXkr8 and hBSG using Fugene○_R_6 (Promega) and cultured for 2 days. Cells were suspended in TBS containing cOmplete EDTA-free, and disrupted by sonication for 30 s at an amplitude of 20 using an ultrasonic disrupter (Q55; Qsonica). Unbroken cells and nuclei were removed by centrifugation at 6,000 × *g* for 25 min, and crude membrane fractions were collected by centrifugation at 99,731 × *g* for 1 h using the Beckman TLA120.2 rotor. Membranes were homogenized in TBS containing 10% glycerol, 1 mM EGTA, and cOmplete EDTA-free, and treated at room temperature for 5 min with 1 mM reduced glutathione. Cells were then incubated at room temperature for 10 min in the presence or absence of copper phenanthroline (0.3 mM CuSO_4_ and 0.9 mM 1,10-phenanthroline). The reaction was stopped by the addition of 0.25 vol. of 200 mM Tris-HCl buffer (pH 6.8), 10% SDS, 25% glycerol, 25 mM N-ethylmaleimide, and 0.05% bromophenol blue. Proteins (0.75 μg) were separated on 10-20% SDS-PAGE, transferred to a PVDF membrane, blotted with HRP-labeled rabbit anti-GFP Ab or HRP-labeled anti-HA Ab (Direct-Blot HRP anti-HA.11 Epitope Tag antibody), and visualized by Immobilon Western Chemiluminescent HRP substrate.

#### Establishment of stable transformants and the phospholipid scrambling assay

PLB and *BSG^-/-^ NPTN^-/-^* W3 cells were transformed by infection with a pantropic retrovirus as previously described^1^. Briefly, HEK293T cells were transfected with a pMXs-puro vector carrying hXkr8-GFP or a pCX4-bsr vector carrying hBSG cDNA tagged with HA or FLAG together with pGP for the gag-pol fusion protein (Takara Bio), and pCMV-VSV-G-RSV-Rev (RIKEN). The virus was concentrated by centrifugation at 6,000 × *g* for 16 h, and used to infect PLB or *BSG^-/-^ NPTN^-/-^* W3 cells. Stable transformants were selected with 1 μg/ml puromycin for hXkr8 and 10 μg/ml blasticidin for hBSG. *TMEM16F^-/-^Xkr8^-/-^* Ba/F3 cells^3^ were transfected with Ahd1-cleaved plasmid DNA by electroporation using NEPA21, and selected with 0.5-1 mg/ml G418 for mBSG and 1.0 µg/ml puromycin for mXkr8 expression. To analyze the cellular localization of Xkr8, stable transformants were observed by confocal microscopy (FV-1000D; Olympus).

Apoptosis was induced by incubating cells at 1.0 × 10^6^/ml with 10 units/ml FasL for 1 h at 37°C as previously described^1^. Phospholipid scrambling activity was measured by the exposure of PtdSer or PtdEtn, or the internalization of NBD-SM^1,21,22^. For the PtdSer-exposure, cells were washed with Annexin V buffer [10 mM Hepes-NaOH buffer (pH 7.4), 140 mM NaCl, and 2.5 mM CaCl_2_], and mixed with 1,000-fold diluted Cy5-Annexin V (for PtdSer) and 5 μg/ml propidium iodide. After an incubation on ice for 15 min, cells were analyzed by flow cytometry with FACSCanto II (BD Biosciences)^1,21^. To incorporate NBD-SM, cells were washed with Annexin V buffer and incubated on ice for indicated time with 0.5 μM NBD-SM at 1.0 × 10^6^ cells/ml in Annexin V buffer. A 150-µl aliquot was mixed with an equal volume of Annexin V buffer containing 5 mg/ml fatty acid-free BSA (Sigma-Aldrich) and 5 nM SYTOX red (Thermo Fisher Scientific), and analyzed by FACSCanto II for the mean fluorescence intensity (MFI)^1^. The PtdEtn-exposure was examined by Ro09-0198-induced cytotoxicity as described previously^4^. In brief, cells at 1.0 x 10^5^/ml were incubated for 15 minutes on ice with Ro09-0198-biotin in Annexin V buffer containing 5 mg/ml BSA. After removing the cells by centrifugation at 300 x *g* for 3 min, the Lactate dehydrogenase (LDH) activity in the supernatant was measured with LDH Cytotoxicity Detection Kit (Takara) according to the manufacturer’s instruction.

## Data availability

The cryo-EM density map for hXkr8-hBSGΔ-Fab18 was deposited in the Electron Microscopy Data Bank (accession number: EMD). The coordinates for the models of the hXkr8-hBSGΔ-Fab18 complex were deposited in the Protein Data Bank (PDB) (accession number). The coordinates and structural factors of Fab14 and the hBSG (domain2)-Fab14 complex were deposited in PDB under accession codes.

## Acknowledgments

We thank Christoph Gerle (Institute for Protein Research, Osaka University) for his advice on EM negative staining and the GraDeR procedure, Jun Suzuki (Kyoto University) for characterizing mouse BSG mutants on N-glycosylation sites, Kazuhiro Abe (Nagoya University) for his advice on the crystallization procedure, the beamline scientists at BL32XU (SPring-8), particularly K. Hirata, for their technical assistance with data collection, F. Takenaga and S. Tsukita (Graduate School of Frontier Biosciences, Osaka University) for their advice on EM negative staining, H. Kawauchi and K. Yamada for their technical assistance, and M. Kamada and M. Fujii for their secretarial assistance. This work was supported in part by Grants-in-Aid from the Japan Society for the Promotion of Science (20K15731 to T.S.; 15H05785 to S.N), Grants-in-Aid from Core Research for Evolutional Science and Technology, Japan Science and Technology Agency (JPMJCR14M4 to S.N.; JPMJCR14M1 to M.K.), and the Platform Project for Supporting Drug Discovery and Life Science Research (Basis for Supporting Innovative Drug Discovery and Life Science Research [BINDS]) of AMED (grant numbers JP17am010172, JP18am0101072, JP19am0101072; support number 0586). This work was also performed in part under the Collaborative Research Program of the Institute for Protein Research, Osaka University (CR-18-05).

## Author contributions

T.S. designed and performed most of the experiments and wrote the manuscript. E.O. helped with protein purification. R.K. H.N. A.N., and C.T. supervised the X-ray crystal analysis and data processing. A.T. and M.K. performed and supervised the cryo-EM analysis. T.M. and T.B. prepared Fab against hBSG. S.N. conceived and designed the project and wrote the manuscript.

---

**Extended Data Fig. 1.**
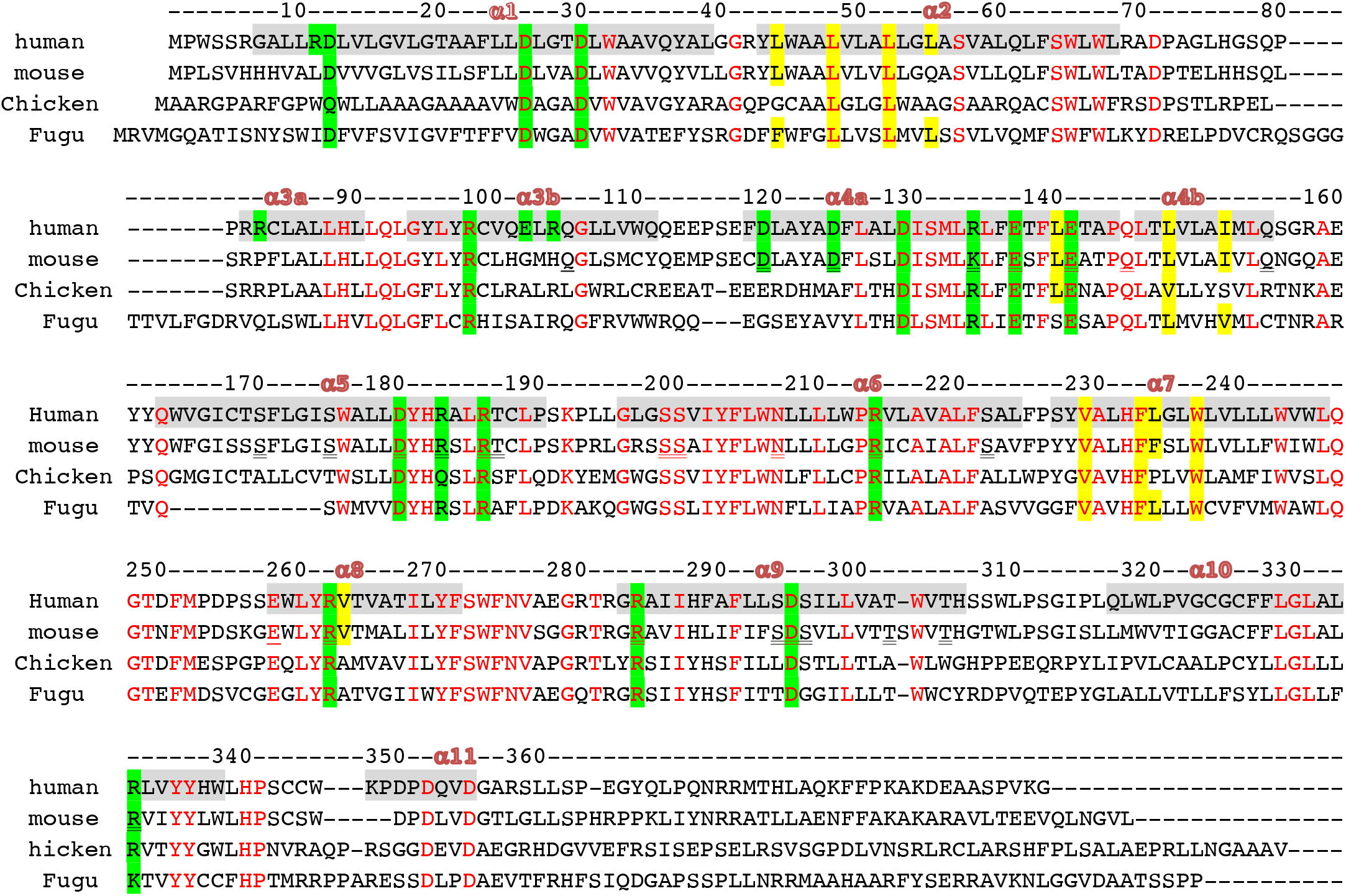
Sequence alignment of Xkr8 orthologues. Sequences of Xkr8 from human (UniProt: Q9H6D3), mouse (UniProt: Q8C0T0), chicken (Chick) (UniProt; Q49M60), frog (UniProt: Q49M63), and fugu (UniProt: H2TYQ9) were analyzed by the MUSCLE Program (EMBL-EBI). Numbers above the first line are the amino acid positions for hXkr8. Conserved residues are indicated in red. Eleven α-helices are shadowed and numbered. Negatively and positively charged residues in the lipid layer are highlighted in green, while the hydrophobic amino acids forming the cleft for PtdCho are highlighted in yellow.

**Extended Data Fig. 2.**
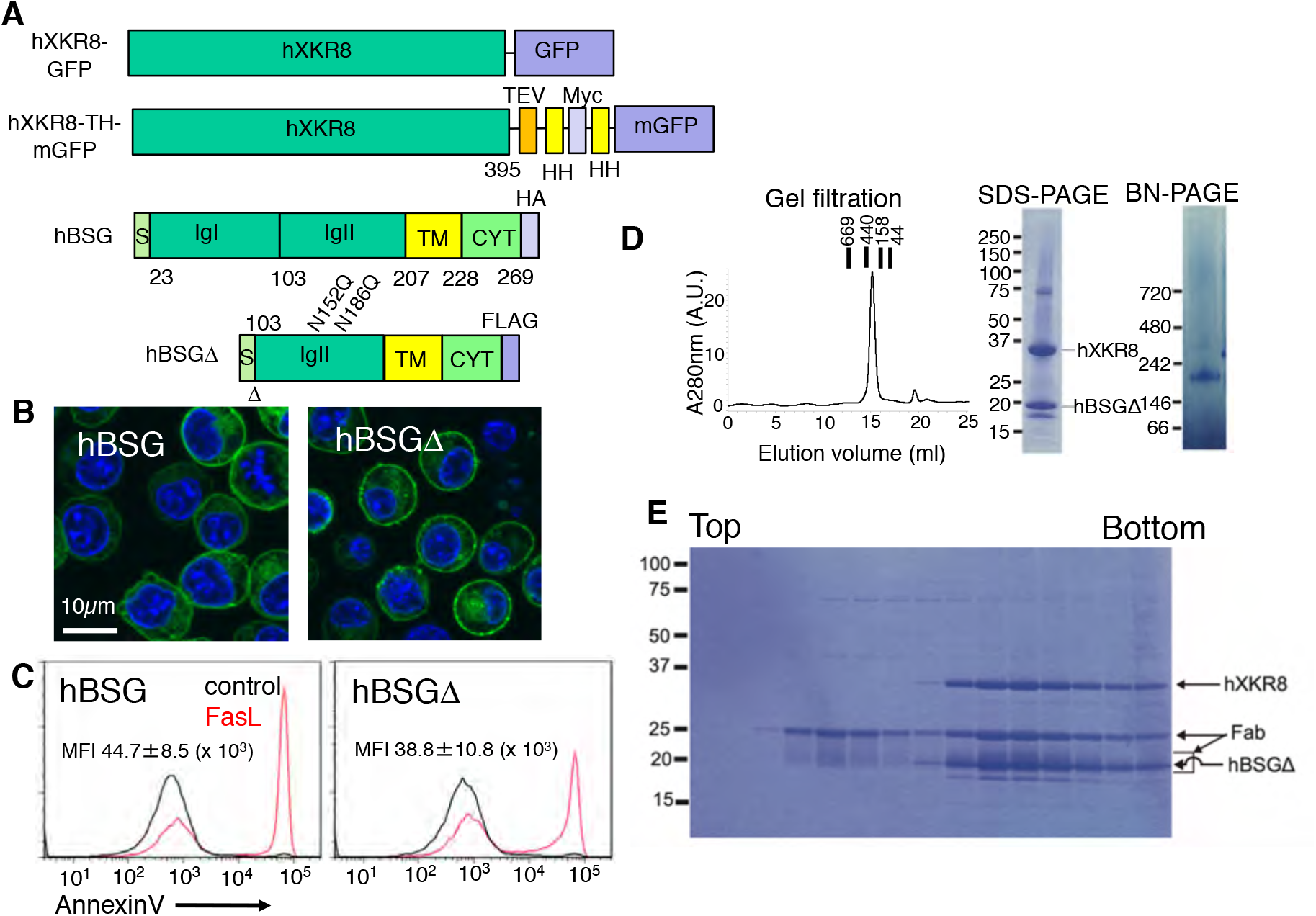
Purification of the hXkr8-hBSGΔ-Fab complex. **a**, Structures of hXkr8-GFP, hXkr8-TH-mGFP, hBSG, and hBSGΔ. mGFP, monomeric EGFP; TEV, TEV cleavage site; HH, Histidine tag (8 His); Myc, Myc-tag; S, Signal sequence; Ig, Immunoglobulin (Ig) domain; TM transmembrane region; Cyt, cytoplasmic region. In hBSGΔ, the first Ig domain (IgI) was deleted, and two N-glycosylation sites (Asn) were mutated to glutamine. **b**, **c** *BSG^-/-^ NPTN^-/-^* W3 cells were transformed by hXkr8-GFP with hBSG or hBSGΔ, and observed by confocal microscopy (**b**). The transformants were treated with FasL, and the Annexin V-staining profile in the PI-negative population was analyzed by flow cytometry (**c**). **d**, The purified hXkr8-hBSG complex was analyzed by gel filtration (7.5 μg protein), SDS-PAGE (6 μg), or BN-PAGE (3 μg). The molecular weight of standard proteins is shown in kDa. **e**, Purified hXkr8-hBSGΔ was incubated with Fab18 and subjected to GraDeR^27^. An aliquot of each fraction was analyzed by SDS-PAGE. hXkr8, hBSGΔ, and Fab (heavy and light chains) are indicated by arrows.

**Extended Data Fig. 3.**
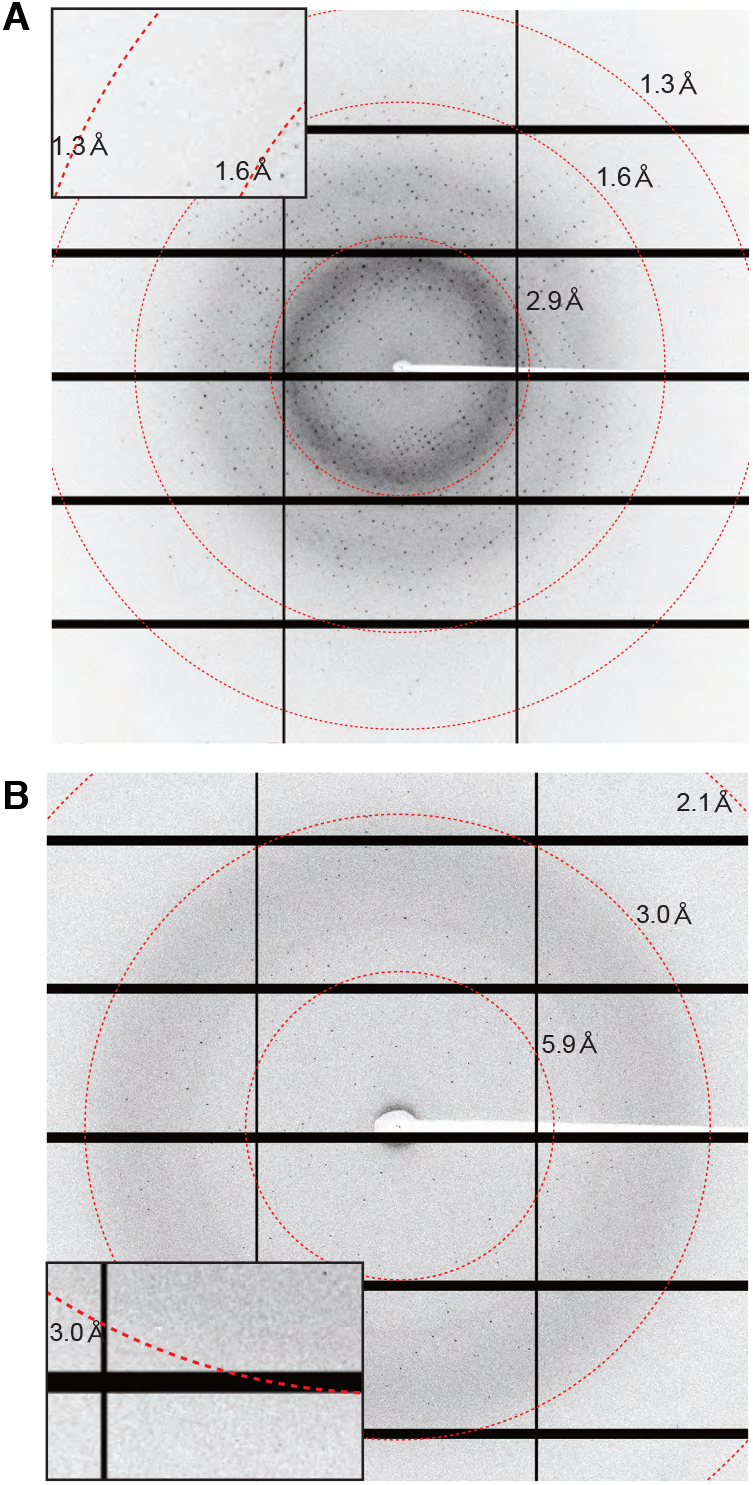
X-ray diffraction analysis of the hXkr8-hBSGΔ-Fab complex. A representative X-ray diffraction pattern of a crystal of Fab14 (**a**) and the lipidated hXkr8-hBSGΔ-Fab complex (**b**) in buffer containing 33% PEG400. The area of high resolution was enlarged in insets.

**Extended Data Fig. 4.**
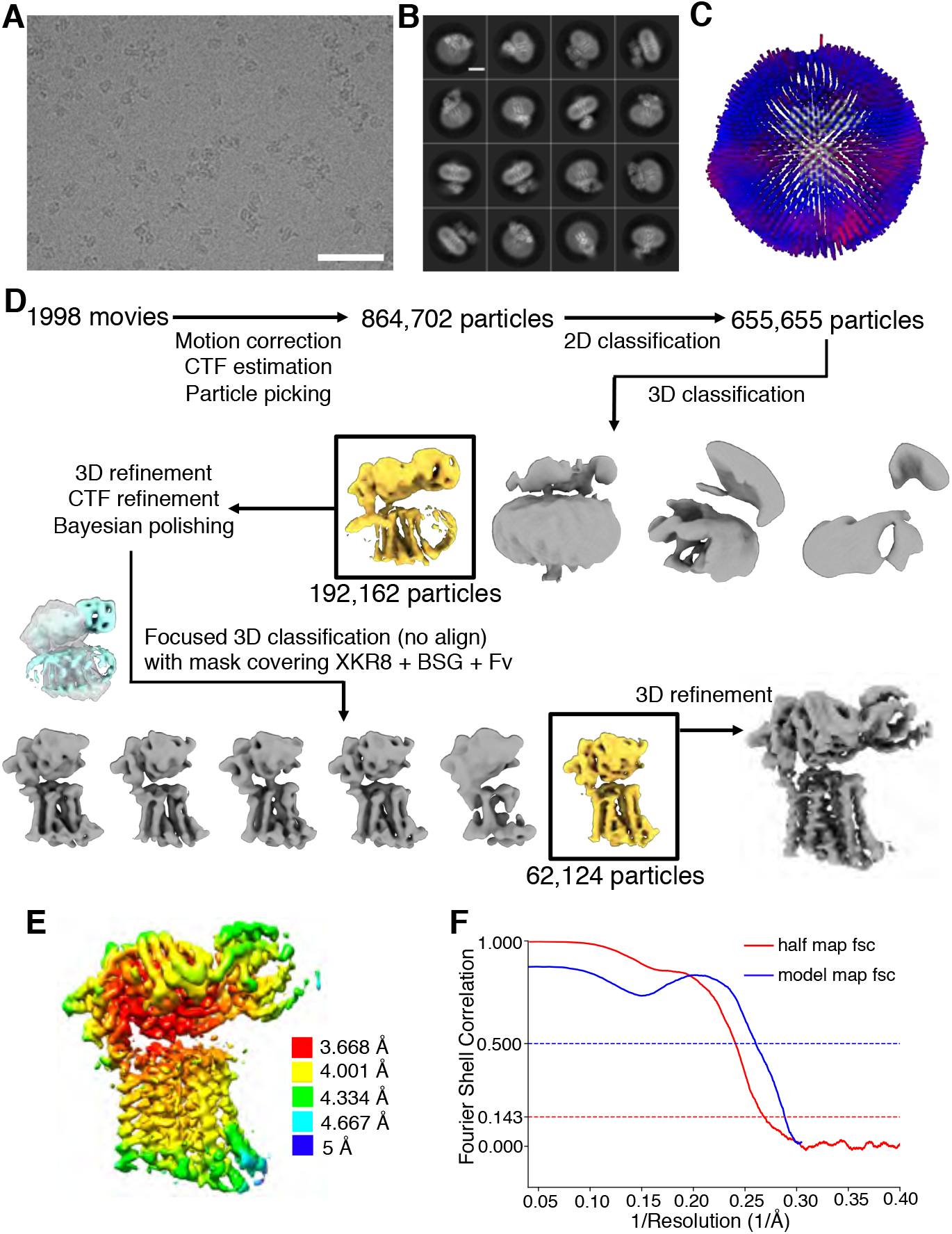
Cryo-EM characterization of the hXkr8-hBSGΔ-Fab complex. **a**, A representative cryo-EM image. Bar, 50 nm. **b**, Representative 2D-class averages, bar, 5 nm. **c**, Angular distribution plot of particles included in the final 3D reconstitution. The number of views at each angular orientation is represented by the length and color of cylinders. Red indicates more views. **d**, Data processing of the hXkr8-hBSGΔ-Fab complex. See Methods for the detailed procedure. **e**, Final reconstruction map colored by local resolution as calculated by RELION3.1. **f**, FSC plot used for resolution estimations and model validation. The Fourier shell correlation (FSC) curve between the EM half maps (red) and FSC curve between the EM full map and atomic model (blue).

**Extended Data Fig. 5.**
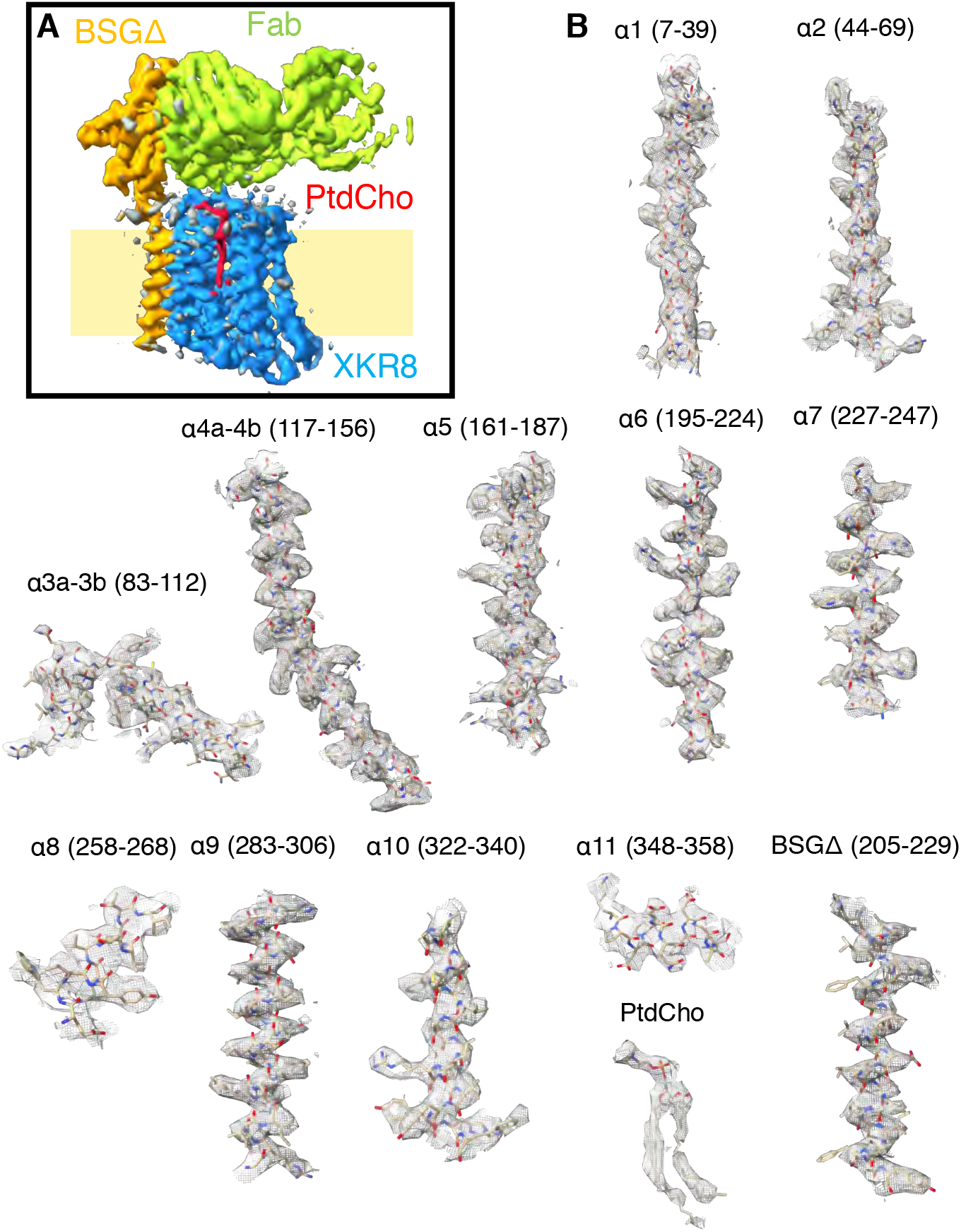
Cryo-EM density of the hXkr8-hBSGΔ-Fab complex. **a**, A cryo-EM map of the hXkr8-hBSGΔ-Fab complex. The location of the membrane is estimated from the position of tryptophan residues (Supplementary Fig. 6). **b**, Representative cryo-EM densities of 11 helices (α1-α11) of hXkr8, the transmembrane helix of hBSGΔ, and PtdCho are superimposed on the corresponding atomic model. Electron microscopy densities are shown in grey meshes, while the model is shown as sticks colored according to the atom type: C, tan; N, blue; O, red; and S, yellow.

**Extended Data Fig. 6.**
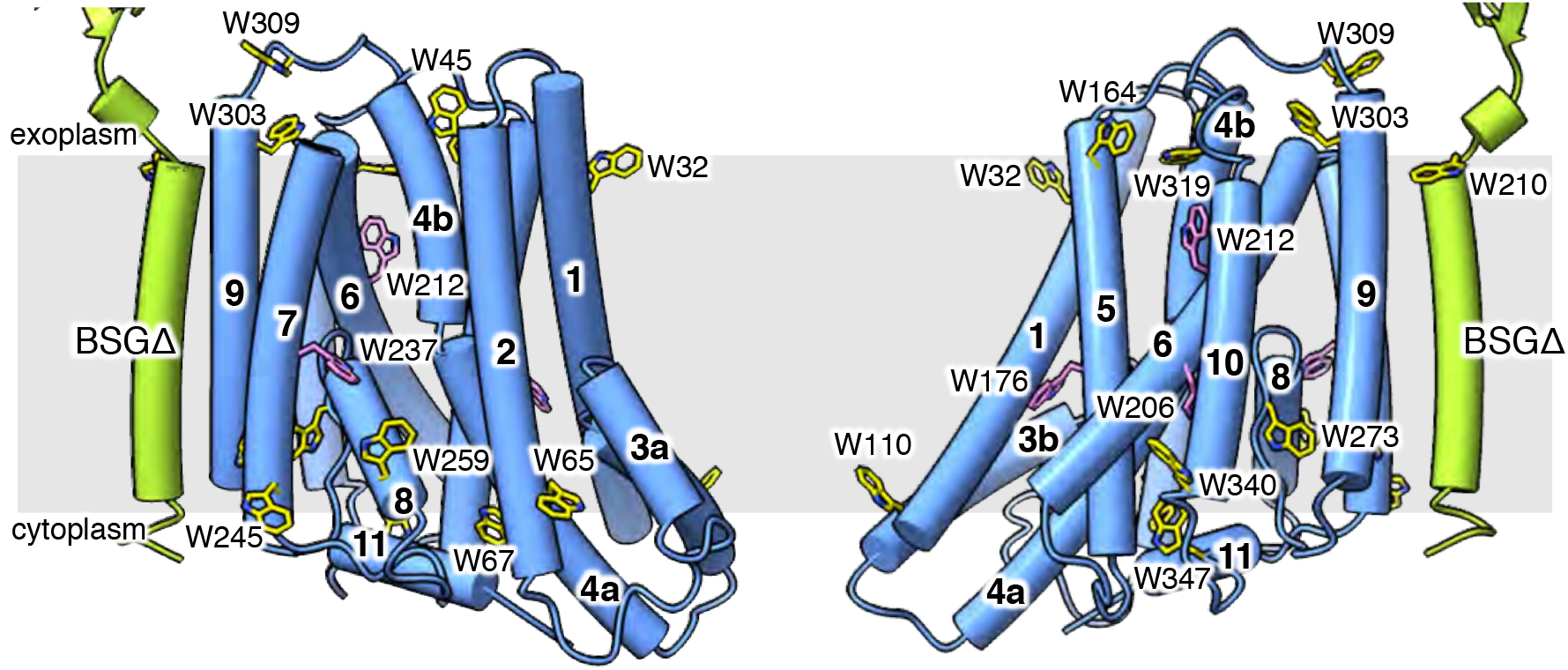
Tryptophan residues in the hXkr8-basigin complex. All tryptophan residues in hXkr8 and the transmembrane region of basigin are shown in the front or rear side views. Tryptophan near the end of the helix is in yellow, while that in the middle of the helix is in magenta.

**Extended Data Fig. 7.**
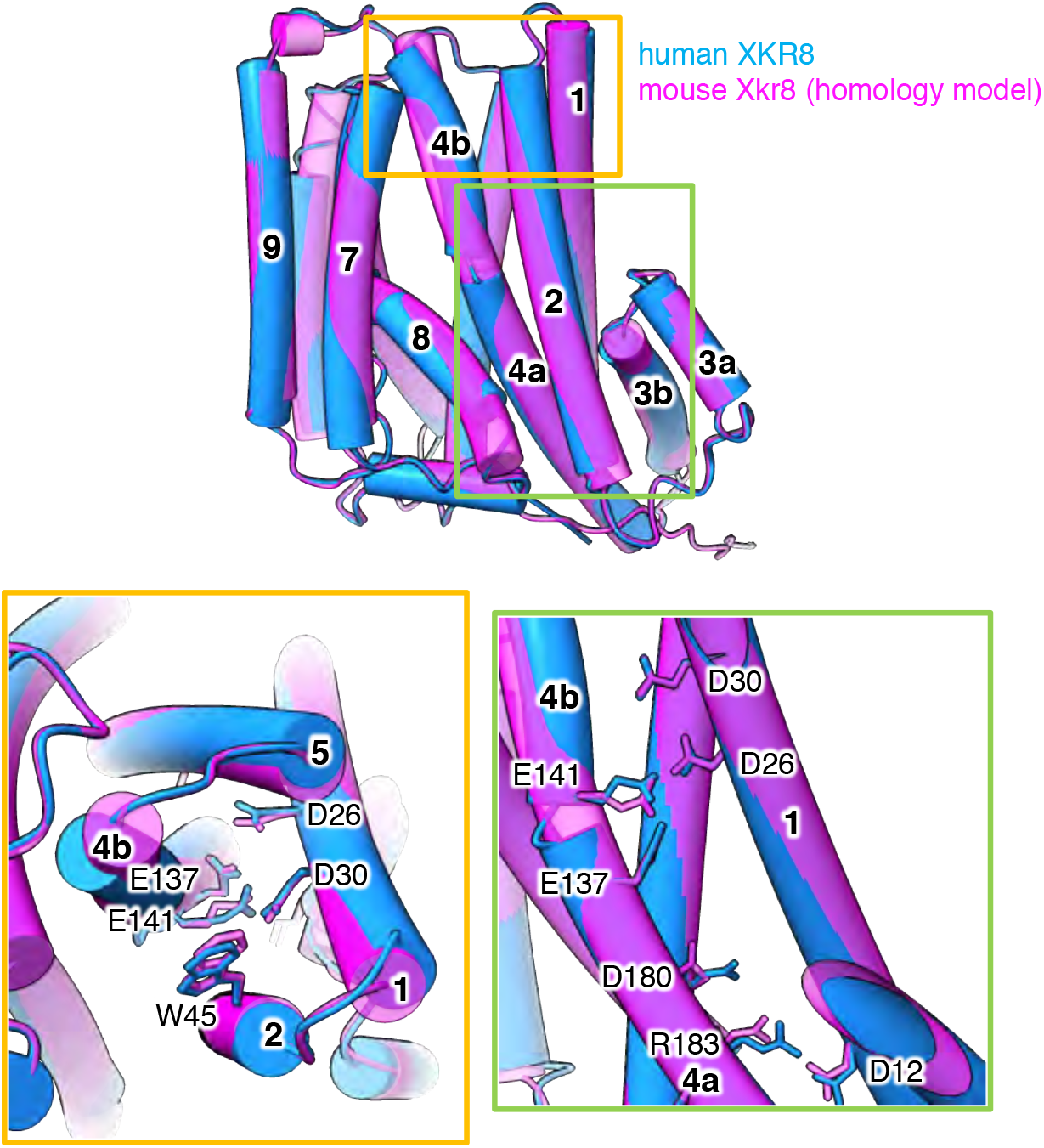
Homology model of mXkr8 structure. mXkr8 homology model was predicted with MODELLER^58^ using the hXkr8 structure (PDB) as a template. mXkr8 homology model is superimposed on hXkr8 structure. Views of the residues involved in stabilizing the structure and those in the phospholipid path are enlarged in the insets.

**Extended Data Fig. 8.**
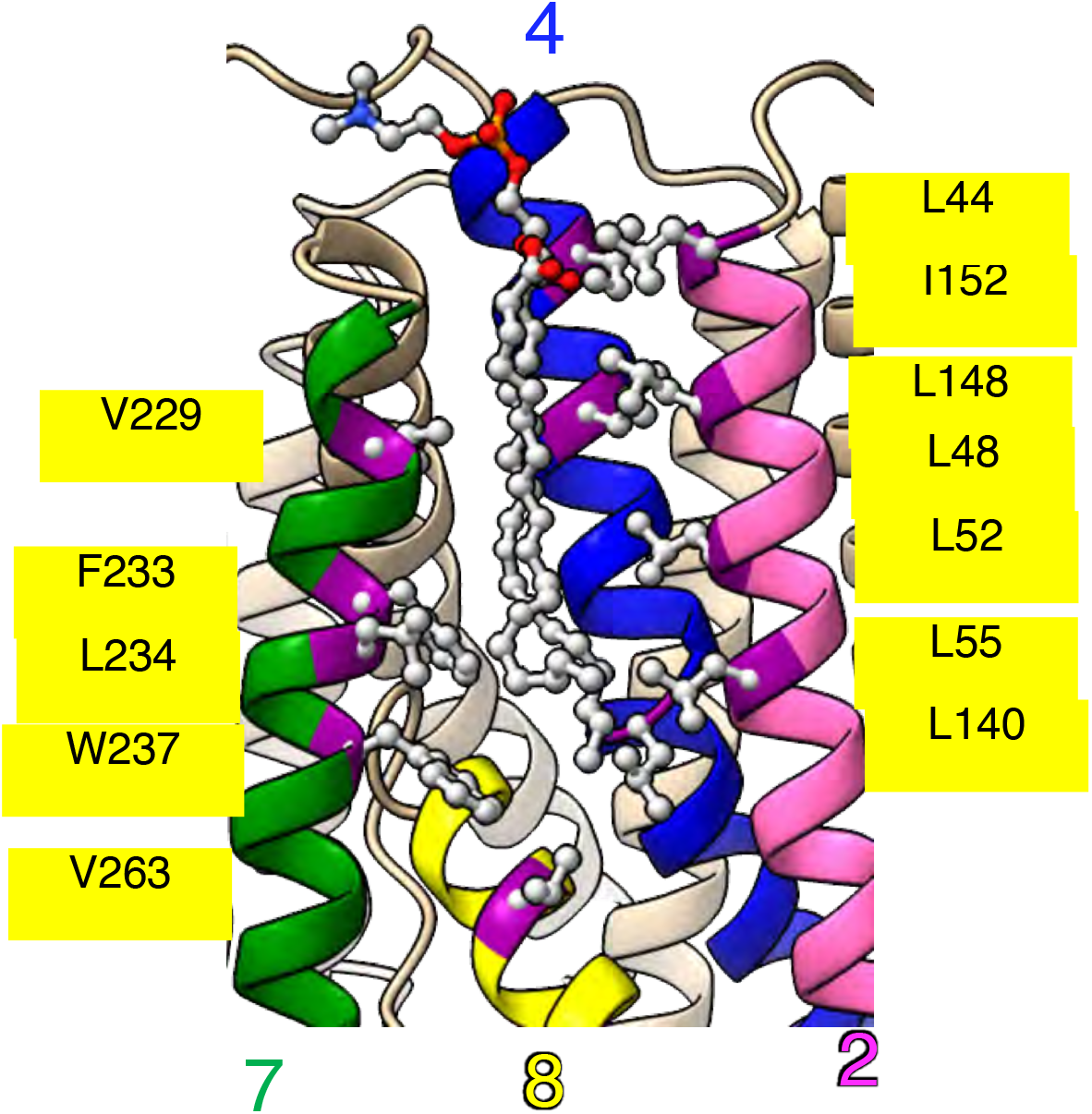
Hydrophobic amino acids surrounding PtdCho. hXkr8 is shown in a ribbon presentation with hydrophobic residues (Leu, Ile, Val, Phe, and Trp) surrounding PtdCho in a ball and stick structure.

**Table S1.** Data collection for X-ray diffraction and refinement statistics.

| Data collection | Fab14 | hBSGext/Fab14 |
| --- | --- | --- |
| Space group | *C*2 | *C*222<sub>1</sub> |
| Cell dimensions |  |  |
| *a*, *b*, *c* (Å) | 163.45, 53.05, 54.31 | 117.35, 247.35, 522.25 |
| $\alpha$, $\beta$, $\gamma$ (°) | 90, 105.05, 90 | 90, 90, 90 |
| Resolution (Å) | 78.924-1.123 (1.194-1.123) | 123.676-2.509 (2.706-2.509) |
| *R*<sub>merge</sub> (all I+ & I-) | 0.048 (0.653) | 0.187 (1.569) |
| *R*<sub>merge</sub> (within I+/I-) | 0.045 (0.659) | 0.178 (1.421) |
| *R*<sub>meas</sub> (all I+ & I-) | 0.054 (0.789) | 0.209 (1.690) |
| *R*<sub>meas</sub> (within I+/I-) | 0.055 (0.897) | 0.220 (1.656) |
| *R*<sub>pim</sub> (all I+ & I-) | 0.024 (0.432) | 0.091 (0.624) |
| *R*<sub>pim</sub> (within I+/I-) | 0.032 (0.604) | 0.127 (0.843) |
| Total number of observations | 668556 (21605) | 18245 (4115) |
| Total number unique | 139276 (6905) | 18245 (571) |
| $\langle I/\sigma(I) \rangle$ | 13.5 (1.4) | 7.3 (1.3) |
| Completeness (spherical) | 81.6 (24.3) | 68.4 (10.8) |
| Completeness (ellipsoidal) | 87.0 (33.7) | 86.3 (45.7) |
| Multiplicity | 4.8 (3.1) | 5.1 (7.2) |
| *CC*<sub>1/2</sub> | 0.999 (0.575) | 0.993 (0.476) |
| **Refinement** |  |  |
| Resolution (Å) | 36.40-1.12 (1.13-1.12) | 123.68-2.51 (2.64-2.51) |
| Number of reflections | 139266 (526) | 18231 (204) |
| *R*<sub>work</sub> / *R*<sub>free</sub> | 0.121 (0.256) / 0.150 (0.333) | 0.228 (0.344) / 0.283 (0.551) |
| Number of non-hydrogen atoms |  |  |
| Protein | 3418 | 3847 |
| Solvent / Ligands / Ions | 695 / 37 / 0 | 0 / 0 / 11 |
| B-factor (Å<sup>2</sup>) |  |  |
| Protein | 19.78 | 38.92 |
| Solvent/ligands/ions | 33.76 / 21.50 / - | - / - / 118.45 |
| r.m.s deviations |  |  |
| Bond lengths (Å) | 0.010 | 0.003 |
| Bond angles (°) | 1.178 | 0.565 |
| Ramachandran (%) |  |  |
| Outlier | 0 | 0 |
| Allowed | 2.63 | 3.57 |
| Favored | 97.37 | 96.43 |

Highest resolution shell statistics are shown in parentheses.

**Table S2.** Cryo-EM data collection and refinement statistics.

|  |  |
| --- | --- |
| **Data collection and processing** |  |
| Magnification | 105,000 |
| Voltage (kV) | 300 |
| Electron exposure (e-/Å<sup>2</sup>) | 48 |
| Defocus range (μm) | 0.8~1.8 |
| Pixel size (Å) | 0.83 |
| Symmetry imposed | C1 |
| Initial particle images (no.) | 864,702 |
| Final particle images (no.) | 62,124 |
| Map resolution (Å) | 3.8 |
| FSC threshold | 0.143 |
| Map resolution range (Å) | 3.67 – 6.51 |
| **Refinement** |  |
| Map sharpening B factor (Å<sup>2</sup>) | -120 |
| Model composition |  |
| Non-hydrogen atoms | 7102 |
| Protein residues | 912 |
| Ligands | 1 (DLP) |
| B-factors (Å<sup>2</sup>) |  |
| Protein | 84.16 |
| Ligand | 66.09 |
| R.m.s. deviations |  |
| Bond lengths (Å) | 0.009 |
| Bond angles (°) | 1.203 |
| Validation |  |
| MolProbity score | 2.45 |
| Clashcore | 11.63 |
| Poor rotamers (%) | 3.27 |
| Ramachandran plot |  |
| Favored (%) | 92.34 |
| Allowed (%) | 7.44 |
| Disallowed (%) | 0.22 |

